# Public good exploitation in natural bacterioplankton communities

**DOI:** 10.1101/2020.12.13.422583

**Authors:** Shaul Pollak, Matti Gralka, Yuya Sato, Julia Schwartzman, Lu Lu, Otto X. Cordero

## Abstract

Microorganisms such as bacteria often interact with their environment through extracellular molecules that increase access to limiting resources. These secretions can act as public goods, creating incentives for exploiters, a.k.a cheaters, to invade and ‘steal’ public goods away from producers. This phenomenon has been studied extensively in microbiology due to its implications for the evolution of cooperation, but little is known about the occurrence and impact of public good exploiters in the environment. Here, we develop a new genomic approach to systematically identify bacteria that can exploit public goods produced during the degradation of polysaccharides. Focusing on chitin – the second most abundant biopolymer on the planet, we show that public good exploiters are active in natural marine microbial communities that assemble on chitin particles, invading during early stages of colonization and potentially hindering degradation. Unlike in classical studies of social evolution, exploiters and polysaccharide degraders are not isogenic and instead belong to distant lineages, facilitating their coexistence. Our approach opens novel avenues to use the wealth of genomic data available to infer ecological roles and interactions among microbes.

## Main Text

In order to access insoluble nutrients in their environment, microorganisms frequently need to invest in costly secretions such as iron-scavenging molecules and hydrolytic enzymes, to name a few (*1*, *2*). This form of extracellular metabolism opens a niche for public good exploiters, or ‘cheaters’, that do not invest in those secretions but reap their benefits. Although many studies have investigated the in-vitro dynamics of otherwise clonal producers and exploiters, the environmental relevance of public good exploitation in a complex environment like the ocean remains poorly constrained. This is largely due to the difficulty of predicting microbial interactions in natural communities. In this paper, we developed a novel approach to predict such interactions in communities that degrade complex forms of organic matter. The microbial degradation of complex organic matter is mediated by extracellular enzymes that break down biopolymers, releasing its fragments to the local environment. Polymer fragments (e.g., oligosaccharides) act as public goods, available as primary carbon and nitrogen sources for all members of the community and not just the enzyme producer. The prevalence of this process leads to the prediction that such communities should be riddled with public good exploiters, or cheaters, which compete directly with degraders by consuming the breakdown byproducts, but do not contribute to the pool of enzymes (*3*). However, community assembly experiments on complex carbon sources have revealed that, although secondary consumers (non-degraders) are numerically dominant, they do not necessarily grow on oligosaccharides and instead prefer to consume metabolic waste products, such as organic acids (*4*, *5*). Therefore, it is unclear whether public good exploitation is relevant in natural biopolymer-degrading communities and if so, how it would contribute to community dynamics and function.

Here, we address this problem in the context of the degradation of chitin, the second most abundant biopolymer on the planet. By detecting genes that are evolutionarily and / or physically linked to chitinases (enzymes that break chitin polymers), we found that exploitation has evolved multiple times across different phyla, suggesting that exploitative lifestyles should be common in natural chitin degrading communities. The study of the genomic signatures associated with chitinases helped us predict which organisms can act as chitooligosaccharide exploiters, without relying on prior protein functional annotation. After validating our predictions, we showed that public good exploiters are present in naturally assembled communities in the ocean, and that their dynamics mirror those of degraders. Finally, we showed that exploiters sampled from natural communities can also stably coexist in the lab with degraders and waste product scavengers, showing that even in these simplified conditions, exploiters do not cause the “tragedy of the commons”, nor are they fully suppressed.

### Predicting exploitation potential from genomes

In theory, detecting a public good exploiter from its genome should be straightforward: an exploiter should lack the genes for public good production (extracellular chitinases, in our case) but have the genetic machinery that allows it to compete for uptake and utilization of the public good against producers. However, as we will show below, predicting which organisms can grow on chitin oligosaccharides in a community is not a simple task. The reason is that competition for oligosaccharide uptake in a community is a complex trait involving multiple processes encoded by different genes (*6*). These genes need not be in the canonical pathway for chitin utilization, as they can mediate surface attachment, biofilm formation, chemotactic behavior, among many other phenotypes. Moreover, given the limitations of gene functional annotations, especially in poorly studied taxa, many of the ecologically relevant genes could lack predicted functions. With this in mind, we first aimed to identify a set of genes that could be used as predictors of the ability of an organism to compete for chitin degradation byproducts in a manner that is independent of functional annotations. We reasoned that if a gene increases the capacity of an organism to grow on chitin oligosaccharides, it should be ‘linked’ to chitinases, either by sharing a similar evolutionary history or by being co-located in the chromosome (*7*, *8*).

We developed a computational approach to detect genes that coevolve or are colocalized with chitinases across 8752 non-redundant complete bacterial genomes spanning the tree of bacterial life (Supplementary Table S1, Supplementary Data S1-2). To infer coevolution, we took advantage of the fact that the number of chitinases in a genome can change drastically even among close relatives, ranging from 0 to 15 chitinases (Fig. 1A), implying frequent gene gain and loss events. By reconstructing the history of gene gain and loss for 3,237,392 gene families across the tree of bacterial life and contrasting these histories against a null model of gene content coevolution (Methods), we found 2097 gene families that coevolve with chitinases in bacteria (Fig. 1B). In addition, we identified 1479 genes that co-localize together with chitinases in the genome, for a total of 3576 candidate predictor genes (Supplementary Data S3). These genes had diverse KEGG annotations such as chemotaxis and motility, attachment and biofilm formation as well as the synthesis of antibiotics, but were most enriched for genes of unknown function and mobile element related functions, highlighting the need for an annotation-agnostic approach (Supplementary Fig. S1, Supplementary Data S4)(*9*). Chitinase-linked genes with nearly identical presence-absence distributions across our species set (e.g., two proteins in the same operon) were further clustered into 1905 “accessory clusters”, containing either individual gene families or small gene family clusters.

**Fig. 1.**
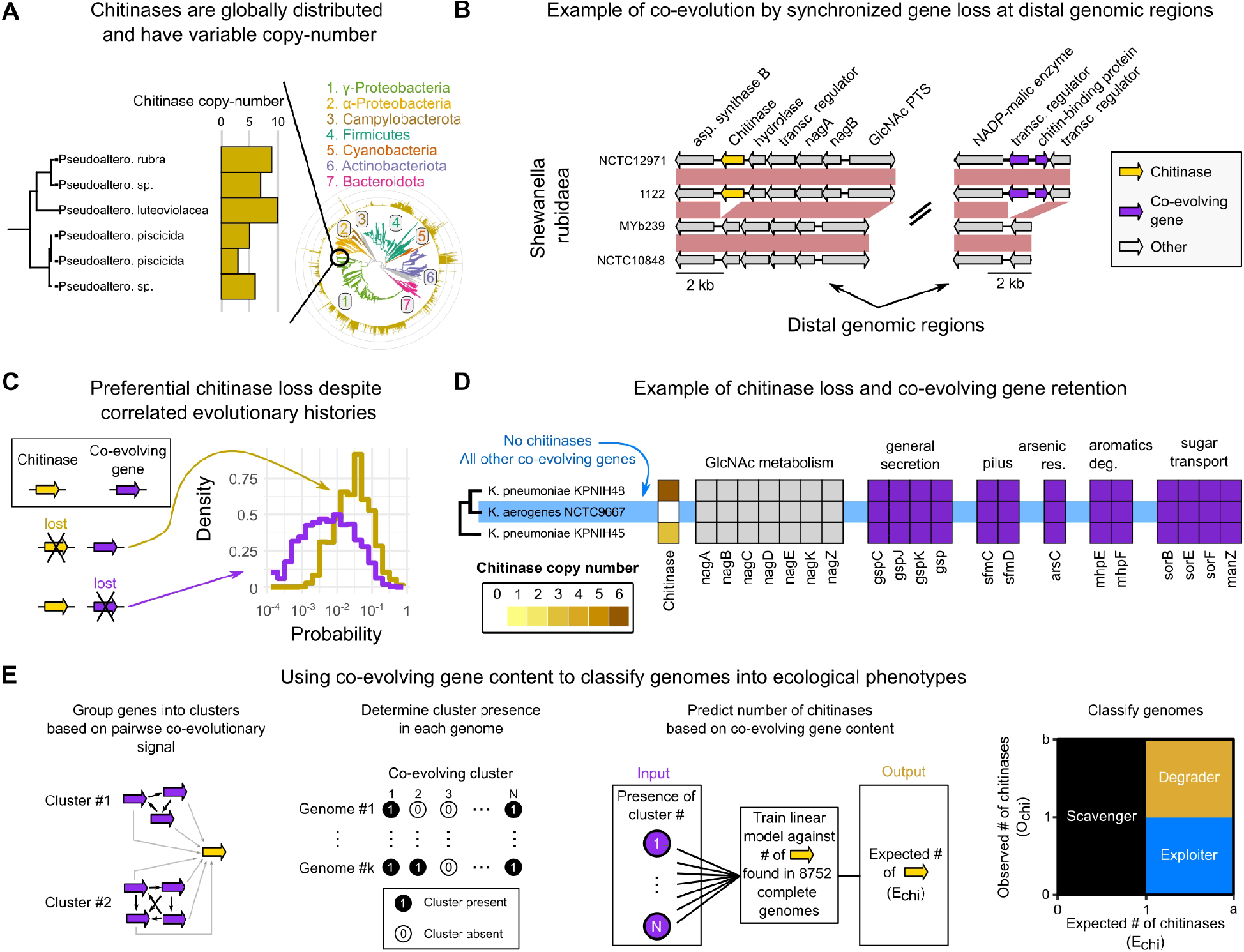
Large-scale detection of genes that co-evolve with chitinases allows gene-content based genome classification. (A) Chitinase copy number in a group of Pseudoalteromonas strains. Inset shows the GTDB-tk derived phylogenetic tree of 8752 genetically distinct (ANI < 99.9%) closed genomes used in this study, with chitinase copy-number indicated on each tip (gold bars). Major phylogenetic groups are highlighted on the tree. (B) An example of coupled gain/loss of a chitinase and two distally located co-evolving genes in a group of closely related Shewanella rubidaea strains. (C) Conditional probability density of losing a chitinase while retaining a coevolving gene (gold) and vice-versa (purple). The probabilities are calculated for each co-evolving gene, and the distribution is over all co-evolving genes. (D) A group of Klebsiella pneumoniae strains, where strain NCTC9667 lost all chitinases while retaining co-evolving genes. Nag genes do not co-evolve with chitinases in this group but are shown to highlight pathway completeness (E) Genome classification from co-evolving gene content. Co-evolving genes are grouped into clusters based on their patterns of co-evolution with each other. Each genome is represented by a bit-string that marks the presence and absence of each cluster. A linear model is trained on all fully sequenced genomes to predict chitinase copy number, given their co-evolving cluster content. New genomes are classified into ecological phenotypes based on the (un)coupling between observed (O_chi_) and expected (E_chi_) number of chitinases.

Although by definition the histories of gain and loss of chitinases and accessory clusters were correlated, chitinases had a higher loss rate than their coevolving genes (mean odds-ratio = 0.755, t-test p-value < 5e-5) (Figure 1C, Supplementary Fig. S2). This means that through evolution there were multiple events in which chitinases were lost but accessory clusters retained, likely leading to the evolution of an oligosaccharide (public good) exploiter. An example is shown in Figure 1D, where a *Klebsiella pneumoniae* strain lost its chitinases, while retaining the genes required for amino-sugar transport and metabolism (*nagABCDEKZ*)(*10*), along with attachment and secretion pili (*sfmCD*, *gsp/gspCJK*)(*11*), as well as other genes involved in arsenic resistance (*arsC*)(*12*), 3-HPP catabolism (*mhpEF*)(*13*), and several sugar transport proteins (*sorBEF, manZ*)(*14*, *15*), all of which coevolve with chitinases. The retention of these genes may be explained by the fact that they are involved in multiple functions, not just chitin degradation. However, independent of the selective pressure that keeps them in the genome, the result of chitinase losses and accessory cluster retention is the evolution of an organism that carries all the genomic machinery needed to compete for chitin oligosaccharides but that does not produce hydrolytic enzymes.

Motivated by these observations, we devised an approach to systematically differentiate degraders, exploiters, and waste product consumers (‘scavengers’), solely on the basis of genomic data. The large list of accessory clusters found to coevolve with chitinases provided us with a set of possible variables to predict an organism’s phenotype, in particular its ability to grow on chitooligosaccharides. Assuming that the number of hydrolases in a genome should correlate with the ability of an organism to grow on the corresponding soluble sugars (*16*), we trained an elastic-net regression model (Methods) to predict the number of chitinases in a genome based on its accessory cluster content. The model was trained separately for each phylum (Methods, Supplementary Table S2) and out of the 1905 coevolving clusters, on average ~190 clusters were sufficient to predict the number of chitinases with high accuracy (average cross-validated R^2^ = 0.825, Supplementary Fig. S3 for comparison with random sets of genes). Interpreting the expected number of chitinases predicted by the model (E_chi_) as a measure of the potential of the organism to compete for chitooligosaccharides, we classified genomes as degraders, exploiters or scavengers based on the contrast between E_chi_ and the actual number of chitinases observed in the genome (O_chi_). If the observed number of chitinases was zero and the expected number was greater or equal to one (O_chi_ = 0, E_chi_ ≥ 1), we predicted the organism to be an exploiter, because it had the genetic machinery that typically accompanies chitinases in genomes of degraders, but lacked the hydrolytic enzyme. If O_chi_ ≥ 1, E_chi_ ≥ 1, we expected the organism to act as a degrader and if E_chi_ < 1 we predicted that the organism was unable to grow on chitooligosaccharides, and therefore must act as waste-scavenger if present in a chitin-degrading community (Fig. 1E).

### Co-evolving gene content successfully predicts mono and co-culture phenotypes

Having trained our model, our next task was to validate its predictions in the context of a naturally assembled chitin-degrading community. To this end, we leveraged a collection of 57 isolates of copiotrophic marine bacteria, co-isolated from the coastal ocean using paramagnetic model marine particles (*4*, *5*). Although our model was not trained on the genomes of this set, we were still able to accurately predict their chitinase content (R^2^ = 0.604). We predicted 13 degraders, 17 exploiters, and 27 scavengers in this collection. Notably, these functional groups were phylogenetically diverse, belonging to such diverse orders as *Flavobacteriales*, *Alteromonadales*, *Rhodobacterales*, and *Vibrionales* (Supplementary Fig. S4). We interpreted our ecological classification (degrader, exploiter, and scavenger) in terms of growth phenotypes that could first be tested using in-vitro mono cultures: degraders should grow on chitin and its monomer GlcNAc, exploiters should grow on GlcNAc but not on chitin, and scavengers should grow on neither of the two substrates (Fig. 2A, Supplementary Fig. S5). We assessed our predictions against those made by genome-scale metabolic models generated using CarveMe, a state-of-the-art tool that derives organism-specific models based on a universal, manually curated set of reactions (*17*).

**Fig. 2.**
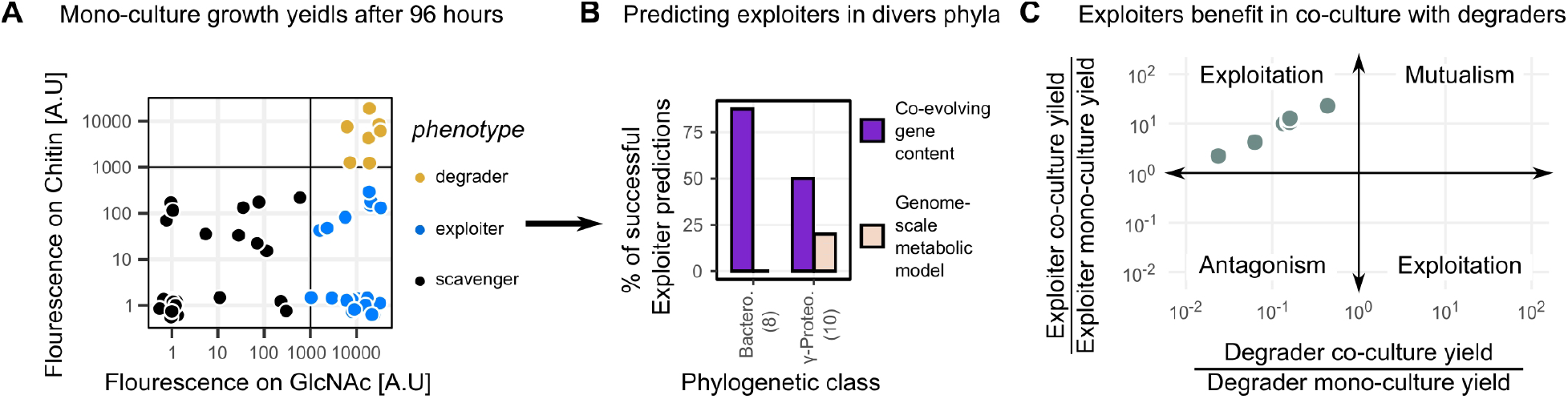
Evolution-guided classification predicts isolate phenotypes in mono and co-culture. (A) Final yield of 57 marine isolates after 96 hours of growth on Chitin and its monomer N-Acetylglucosamine. Yield was measured as fluorescence emitted from incorporated Sybr-gold DNA dye (methods). Each point represents the mean of at least 3 biological repeats. Points are colored according to their observed phenotype. (B) Agreement between predictions made using different genome-informed methods, and actual observed phenotypes. Phenotypically, exploiters are isolates that grow on GlcNAc (>1000 a.u), but not on chitin (<1000 a.u) (C) Co-cultures between 2 degraders and 3 *Flavobacteria* exploiters. Exploiters were misclassified by the genome-scale metabolic model, but correctly classified by their co-evolving gene content. Growth was conducted for 72 hours in minimal media with chitin as the sole carbon source. The x-axis depicts the fold change in degrader yield during co-culture compared to monoculture, and the y-axis shows the same quantity for the exploiter. Each point represents a different co-culture, and represents the mean fold-change from 2 independent replicates. Raw data is for the co-cultures is presented in Supplementary Table S4.

Overall, our model performed better than the annotation-based predictions, making ~30% fewer errors (18 vs 26 errors, Matthews correlation = 0.51 vs 0.28), confirming the value of our approach (Supplementary Fig. S6). Critically, virtually all of the improvement came from the ability of our model to correctly predict exploiters, which the genome-scale metabolic model misclassified as scavengers (Fig. 2B, Supplementary Table S3). In particular, we find that the gap-filled genome-scale metabolic model underperforms when predicting the phenotypes of two relatively understudied families, *Flavobacteriaceae* and *Alteromonadaceae* (Fig 2B, Supplementary Fig. S7). This is consistent with the notion that models that rely on functional annotations should perform poorly on organisms that are distant from the few that have been experimentally characterized, such as *E. coli*, *B. subtilis*, or *V. cholerae*. In contrast, our approach only requires sampling enough genomic diversity and a single annotation (in this case, chitinases), and can therefore be applied to any uncultured and poorly annotated organism.

Ultimately our predictions make a statement about interactions between co-cultured community members. Plasmid encoded fluorescent reporters allowed us to study the dynamics of a *vibrio1A01* (*Vibrio*) and *alteroA3R04* (*Alteromonas*) co-culture in detail, when these two strains were grown with chitin as the sole carbon source. This experiment showed that the exploiter (*alteroA3R04*) was only able to grow in the presence of the degrader, at the cost of the degrader’s final yield but not its maximal growth rate (Supplementary Fig. S8-10). This result prompted us to examine more pairs to confirm our ability to predict exploitative interactions based on the final yields of the co-cultured members. To this end, we co-cultured several degrader-exploiter pairs in minimal-media containing chitin as the sole carbon source for 72 hours. We used 2 different degraders and 3 isolates that were predicted by our method to be exploiters, but were classified as scavengers by the genome-scale metabolic model. We found that, as predicted, in all cases the interaction was exploitative: putative exploiters produced on average ~10-fold more colony forming units (CFUs) compared to growth in monoculture. Degraders, on the other hand, suffered from the presence of exploiters, decreasing their yield by a factor of ~6 on average (Fig. 2C). Interestingly, we found that in those cases where degraders were suppressed by cheating, the exploiter did not reach a high yield, and vice-versa. This trade-off between exploitation and degrader yield can be recapitulated with simple consumer-resource models that include public good exploitation (Supplementary Fig. S12), lending further support to the notion that our method is able to successfully predict GlcNAc consumers (i.e. exploiters) where annotation-based methods fail.

### Dynamics of exploiters in the environment

Our in-vitro results do not guarantee that our putative exploiters can invade natural chitin-degrading communities in a manner consistent with their purported role. To address this question, we studied the population dynamics of our isolates in their native seawater community. We mapped the isolates’ 16S rRNA sequences to the time-series data of the original enrichment experiment, in which the community assembled from seawater onto chitin particles for a period of 244 hours (Fig. 3A)(*4*, *5*). This allowed us to analyze the colonization dynamics of each mapped isolate across the different ecological roles (Fig. 3B). We found that the colonization dynamics of exploiters were more similar to that of degraders than to that of scavengers, consistent with their role as parasites of degraders. Exploiters reached maximum frequency in the succession shortly after degraders (12-16 h for exploiters, 8 h for degraders), while scavengers peaked at late successional stages (>80 h) (Supplementary Fig. S11). This observation implies that the accessory gene clusters, containing the genetic machinery to chemotax towards GlcNAc, adhere to chitin surfaces, etc., are the main determinant of early colonization, and not chitinases themselves. In other words, our response variable, E_chi_, should be a better predictor of colonization dynamics in a wild community than chitinase copy number (O_chi_). To test this assertion, we calculated the maximal speed of colonization of each isolate during the early phases of assembly, before the community became dominated by scavengers (Methods, Fig. 3C). As expected, we found that E_chi_ was a better predictor of colonization speed than the observed number of chitinases, O_chi_. Genomes predicted to contain chitinases had a 19-fold higher colonization rate on average compared to those predicted to lack chitinases (p-value = 0.02). In contrast, genomes encoding chitinases did not display a significant difference in colonization rate compared to those that lacked chitinases (p-value = 0.37, average fold-difference = 2.1).

**Fig. 3.**
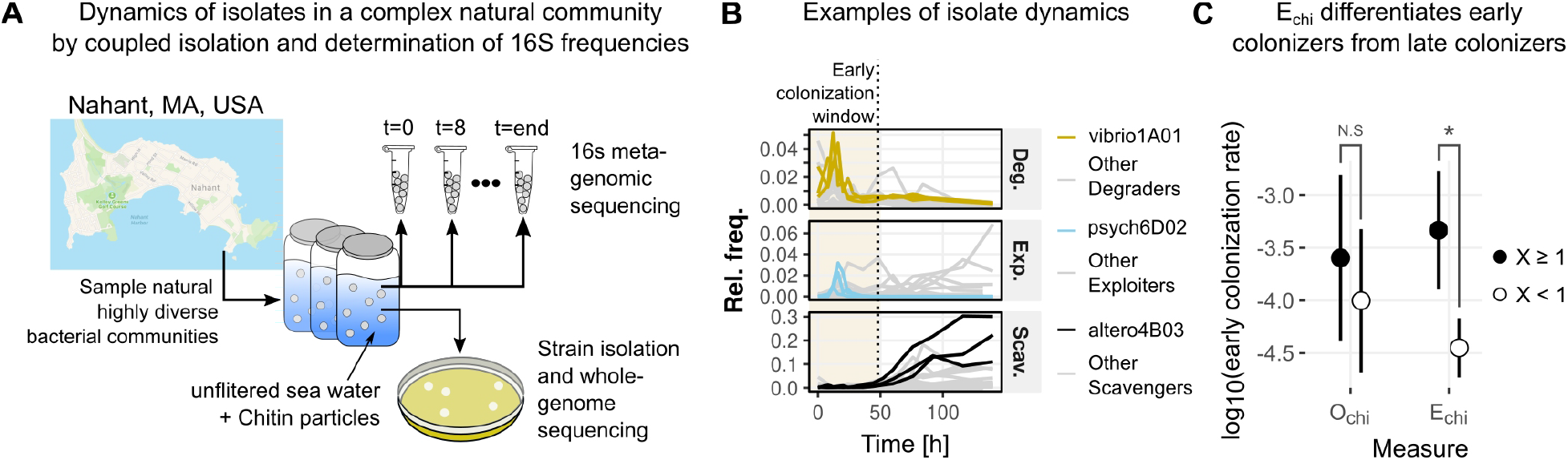
Exploiter dynamics mimic those of degraders in complex natural communities. (A) Natural, highly diverse bacterial communities were sampled as sea water samples taken from Nahant, MA, USA, and incubated with chitin as the sole carbon source. Samples from the incubation were taken periodically for DNA extraction and isolate collection. (B) Colonization dynamics of 16S sequences that perfectly matched the 16S sequences of isolated strains. Isolates were assigned an ecological role according to their co-evolving gene content. Each panel shows the trajectories of a different ecological role, and each line represents the trajectory of a single 16S sequence in a single repeat. 1 representative strain from each role is highlighted as indicated by the legend. The early colonization window was defined as in (*4*) (C) Grouping strains by the expected number of chitinases (E_chi_), and not by their observed number (O_chi_), results in a clean separation between early colonizers (degraders and exploiters) and late colonizers (scavengers). Asterisk marks p-value < 0.05. N.S = p-value > 0.3).

### Exploiters, degraders and waste scavengers coexist in synthetic communities

In the natural chitin-degrading communities the dynamics of colonization and growth are successional, meaning stable coexistence cannot be attained on a single chitin particle. Instead, degraders, exploiters and scavengers could stably coexist at the larger scale of many particles (the metapopulation). To test whether the degraders and exploiters could coexist in a closed system with chitin particles as the sole carbon source – or whether their overlap in resource preference would lead to an eventual community collapse, we assembled synthetic communities with 44 strains sampled from our isolate collection and from the three different roles. The strains were chosen based on 16S sequence distance and robust preculture growth. To assess coexistence, we serially passaged the co-culture over 11 dilution-growth cycles (Fig. 4A,B, Supplementary Data S5) and repeated these experiments with three different dilution factors (which determine the minimum growth rate required to survive serial passages) to assess the robustness of coexistence to different growth conditions (Methods).

**Fig. 4.**
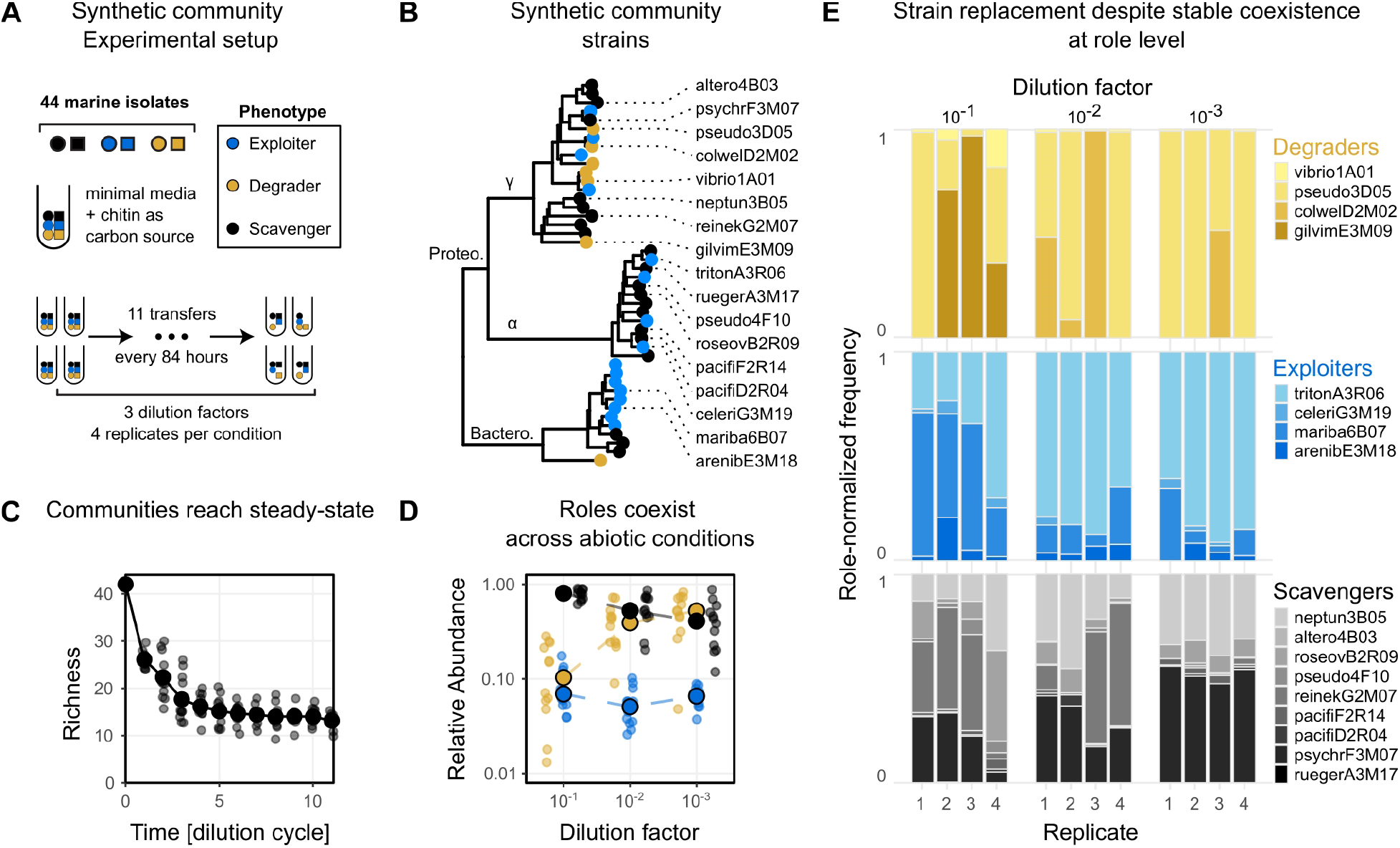
Synthetic communities display stable coexistence of ecological roles, with stochastic strain replacement. (A) Synthetic communities were assembled from 44 isolates that represent different phylogenetic groups and ecological types (Supplementary Data S5). The initial composition was identical for all communities, as they were sampled from the same initial mix. Communities were allowed to grow for 84 hours and were transferred 11 times. We performed 3 different dilution factors (10^-1^, 10^-2^, and 10^-3^), with 4 repeats per condition. (B) GTDB-tk generated phylogenetic tree of the isolates used in this experiment. Isolate phenotypes are indicated by colored circles at the tip of the tree. Isolates that survived to the end of the experiment in at least one condition are indicated by name. (C) The number of species present in a community (Richness) as a function of time (measured in number of growth-dilution cycles). (D) Steady-state relative abundance (transfer > 8) of each ecological role in the synthetic communities. Each point represents the total role abundance from a given sample at a specific time point. Large circles represent the medians for each role as a function of the dilution factor. Colors represent ecological roles as in panel A. (E) Each bar represents the role-normalized abundance of all isolates belonging to a certain ecological role at the final time point. After normalization, abundances of strains with same role add up to 1.

Following a short transient where species are rapidly purged, community richness stabilized on 10-15 members (Fig. 4C), with coexistence of all three roles across all dilution factors (Fig. 4D). Over the final five transfers, after community richness had equilibrated, exploiters were stably maintained but at low abundance, recruiting roughly 10% of all reads. Scavengers, by contrast, were the most abundant ecological group (20-80%), supporting previous findings of widespread export of metabolic waste by exploiters and degraders (*4*, *5*). Scavenger and degrader abundances were controlled by the dilution factor, with degraders increasing in relative abundance with increasing dilution strength (Fig. 4D). This trend can be interpreted as a consequence of the lower time-averaged growth rates of scavengers, which have to wait for the accumulation of secreted metabolites to grow.

It is evident that replicates can be variable in terms of the frequencies of the different ecological roles (Fig 4D). Closer inspection of the community compositions across dilution factors and replicates revealed different patterns of diversity within ecological roles. In most communities, degraders were dominated by a single strain*, pseudo3D05* (*Pseudoalteromonas*), and two exploiters, *mariba6B07* (*Maribacter*) and *tritonA3R06* (*Tritonibacter*), while four to five scavengers coexisted in the same community. However, there were clear cases in which other strains became dominant: *pseudo3D05* coexisted with other degraders at appreciable abundance in at least four out of the twelve communities; was altogether replaced by *colwelD2M02* (*Colwellia*) in one replicate and *gilvimE3M09* (*Gilvimarinus*) in another, while one of the two exploiters seemed to reach higher relative abundances at higher dilution factors. The abundance of different scavengers varied across replicates in a seemingly stochastic manner, especially at low dilution factors (Fig. 4E). These compositional changes within the different roles seemed to be independent of each other, i.e., the different strains were substituted with no apparent consequence for the rest of the community. The fact that the roles stably coexisted despite variation in community composition suggests that degraders, exploiters, and scavengers represent three stable metabolic niches in polysaccharide degrading communities.

## Discussion

In this study we developed a novel computational approach that allowed us to predict phenotype from genome, in particular the ability of an organism to act as a chitologosaccharide consumer. A key feature of our method is that it is agnostic to functional annotations and instead builds on evolutionary patterns that can be inferred directly from genomic data. The fact that genome repositories continue to grow at an accelerated pace, whereas functional annotations remain limited to what can be learned from a few model organisms, underscores the relevance and timeliness of our approach. Ancestral genome reconstructions have been used for many years to infer gene-gene coevolution, with the goal to expand and guide the discovery of protein-protein interactions in microorganisms (*18*). However, the work presented in this paper is the first to leverage these methods to infer and validate the growth phenotypes of microbes and their ecology in complex communities.

Using this approach, we showed that oligosaccharide exploiters, which share many genomic features with degraders but do not encode the relevant hydrolytic enzymes, are abundant in natural communities and have the potential to hinder degradation. The frequent losses of chitinases during the evolution of bacteria are likely the result of short-term evolutionary incentives created by the release of public goods during extracellular hydrolysis. For degraders, in turn, the invasion of exploiters can create incentives to privatize the public good, via spatial aggregation or tethering of enzymes to the membrane (*19*). Recent studies have also shown that some *Flavobacteria* are capable of a ‘selfish’ mode of uptake, whereby long oligomers are directly incorporated inside the cell, bypassing the need for extracellular hydrolysis (*20*). The relative advantage of one strategy over the other and the conditions where each may be more relevant, remain still unknown. However, what is clear is that the constant tension between degraders and exploiters seems to have shaped the evolution of polysaccharide degradation strategies among bacteria, highlighting the fact that microbial interactions and the evolutionary dynamics they generate can impact key ecosystem processes, such as the degradation of organic matter.

Although consistent with the notion of a social ‘cheater’, there is an important difference between the exploiters we identified and cheaters, as typically conceived in the social evolution literature: cheaters are loss-of-function mutants that rise in frequency due to a short-term fitness advantage over a wild-type cooperator phenotype (*21*). Natural exploiters of polysaccharide degraders, by contrast, invade communities by a process of community assembly. Consequently, exploiters in naturally assembled chitin-degrading communities are distantly related to degraders and have diverse metabolic potentials. This increases the possibilities for coexistence, since organisms belonging to these two roles can differ along multiple phenotypic dimensions, but it also makes it harder to predict interspecific interactions. Our in-vitro studies indicate that, under simple batch culture conditions, public good exploiters hinder the growth of degraders, potentially slowing down the turnover of organic matter. However, further work is needed to elucidate the impact of this common ecological interaction in more realistic conditions, such as under fluid flow and multiple nutrient limitations.

## Acknowledgments

We would like to thank Martin Polz, Seppe Kuehn, Avigdor Eldar, Akshit Goyal, and members of the Cordero lab for their helpful comments.

## Author contributions

Shaul Pollak: Conceptualization, Investigation, Methodology, Visualization, Writing. Matti Gralka: Investigation, Visualization, Writing. Yuya Sato: Methodology, Julia Schwartzman: Methodology. Otto X. Cordero: Conceptualization, Project admin, Funding, Surpervision, Writing.

## Data and materials availability

All data is available in the main text or the supplementary materials. Code available at https://github.com/sigmap666/NaturalExploiters

