## supplemental information for "Public good exploitation in natural bacterioplankton communities"

**This PDF file includes:**

Materials and Methods
Figs. S1 to S12
Tables S1 to S4
Captions for Data S1 to S5

**Other Supplementary Materials for this manuscript include the following:**

Data S1 to S5

### Materials and Methods

Strains, plasmids, and growth conditions

Marine-broth (MB) was purchased from Millipore-Sigma (catalogue #76448). The minimal media used throughout (MBL) was prepared by mixing 10-2 [M] NH4Cl, 10-3 [M] Na2HPO4, 10-3 [M] Na2SO4, 0.05 [M] HEPES (pH 8.2), 20 [g/L] NaCl, 3 [g/L] MgCl2*6H2O, 0.15 [g/L] CaCl2*2H2O, and 0.5 [g/L] KCl. The following trace metals and vitamin solutions were added at a final concentration which is 1/1000 the indicated concentrations. Trace metals: 2100 [mg/L] FeSO4*7H2O , 30 [mg/L] H3BO3, 100 [mg/L] MnCl2*4H2O, 190 [mg/L] CoCl2*6H2O, 24 [mg/L] NiCl2*6H2O, 2 [mg/L] CuCl2*2H2O, 144 [mg/L] ZnSO4*7H2O, 36 [mg/L] Na2MoO4*2H2O, 25 [mg/L] NaVO3, 25 [mg/L] NaWO4*2H2O, 6 [mg/L] Na2SeO3*5H2O. Vitamins (dissolved in 10 mM MOPS pH 7.2): 100 [mg/L] Riboflavin, 30 [mg/L] D-Biotin, 100 [mg/L] Thiamine hydrochloride, 100 [mg/L] L-ascorbic acid, 100 [mg/L] Ca-d-pantothenate, 100 [mg/L] Folate, 100 [mg/L] Nicotinate, 100 [mg/L] 4-aminobenzoic acid, 100 [mg/L] pyridoxine HCl, 100 [mg/L] Lipoic acid, 100 [mg/L] NAD, 100 [mg/L] Thiamin pyrophosphate, 10 [mg/L] Cyanocobalamin.

Strains, their phylogenetic placement, and inferred ecological role are provided in Data S3. Plasmid pVSV208 was a kind gift from the Ruby lab (*1*). It contains an origin of replication from the plasmid pES213, the R6K origin of replication, an origin of transfer, and encodes chloramphenicol resistance and the fluorescent protein DsRed. Plasmid pVSV208 was maintained in Escherichia coli strain DH5-alpha lambda pir or PIR1 (Invitrogen). The strain was routinely grown in LB with 25 µg/mL chloramphenicol. Plasmid pBTK569 was a kind gift from the Barrick lab, and was acquired from Addgene (Addgene plasmid #110614; http://n2t.net/addgene:110614; RRID:Addgene_110614). It contains the RSF1010 origin of replication, an origin of transfer, and encodes spectinomycin resistance and the fluorescent protein mCrimson. Plasmid pBTK569 and its derivative pLL014 were maintained in E. coli Top10 (Invitrogen). 50 µg/mL spectinomycin was used to select for pBTK569, and 25 µg/mL chloramphenicol was used to select for pLL014. *E. coli* strain DH5 alpha was used to maintain pEVS104 and was grown in LB containing 25 µg/mL kanamycin.

To construct pLL014, a variant of pBTK569 encoding chloramphenicol resistance and eGFP, we synthesized a segment of DNA coding for eGFP, and the chloramphenicol-resistance gene cat (GeneArt, Invitrogen). We amplified this segment with primers engineered with the restriction sites BsrGI and XbaI, which are present at unique sites in pBTK569, using high-fidelity Q5 polymerase (NEB). We digested pBTK569 and the Cm-eGFP DNA with BsrGI and XbaI (NEB) and treated the plasmid digest with Antarctic phosphatase to prevent re-ligation. We ligated the digested Cm-eGFP and pBTK569 with T4 DNA ligase (NEB), and electroporated the products into Top10 *E. coli* (Invitrogen). The construct was confirmed by Sanger sequencing.

Plasmids pLL014 and pVSV208 were introduced into strains of interest by conjugation. The helper plasmid pEVS104 (*2*), which carries in-trans copies of the conjugative transfer genes, was used to mobilize the oriT containing plasmids. Equal parts of overnight cultures of E. coli strains carrying pVSV208 or pLL014 (donor), pEVS104, or the recipient strain were washed, resuspended in antibiotic free MB, combined, and pelleted. The pellet was resuspended in a small amount of MB, and plated on MB 1.5% agar. After overnight incubation at 25 °C, the conjugation mixture was plated onto MB-agar containing 12.5 µg/mL chloramphenicol. Colonies were plated on two rounds of 12.5 µg/mL MB agar to isolate plasmid-carrying recipient strains from any residual E. coli. The transconjugants were confirmed by sequencing the 16s rRNA gene using universal primers 8F (5-AGAGTTTGATCCTGGCTCAG-3) and 1522R (5-AAGGAGGTGATCCANCCRCA-3).

For isolate phenotyping (Figure 2A), strains were streaked from glycerol frozen stocks onto MB-agar plates and allowed to grow for 48 hours at 25˚C. Single colonies were inoculated into 14 [ml] Polypropylene growth tubes (VWR #60818) containing 1 [ml] MBL media supplemented with either 20 [mM] pyruvate or 20 [mM] glucose and grown for a maximal duration of 48 hours at 25˚C with shaking. Whenever possible, cultures grown for 24 hours in MBL + 20 [mM] pyruvate were used. If a strain did not show visible growth, as measured by optical density, after 24H in MBL + 20 [mM] pyruvate, the MBL + 20 [mM] glucose was assayed for optical density. If the glucose culture also showed no visible growth, this strain's culture was allowed to grow for an additional 24H, and the cultures were checked in the same order again. After pre-growth, cells were washed three times in artificial seawater (Millipore-Sigma #S9883) by centrifugation at 5000 rpm, and diluted 1/20 into the final growth media. Assays were performed in (biological) triplicate in deep 96-well plates containing 1 ml of MBL media supplemented with either 20 [mM] GlcNAc or 2 [g/l] colloidal chitin. The same protocol was used for the final C.F.U yield measurements of co-cultures as presented in Figure 2C.

For the co-culture dynamics experiments (Supplementary Figure S11) a predicted exploiter candidate, alteroA3R04 labeled with a plasmid-encoded eGFP (plasmid pLL014), and a chitin degrader, vibrio1A01 labeled with a plasmid-encoded DsRed (plasmid pVSV208), were co-cultured with a chitin sheet as a sole carbon source. The transparent chitin film was generated at the bottom of the well of a 96-well plate by adding 32 µl of 10 mg/ml chitin solution (dissolved in HFIP) and allowing the solvent to dry. The cells of the two strains were pre-cultured in MBL + 20 [mM] GlcNAc at 25°C, and diluted using MBL salts to an OD600 of 0.0625. The two strains were then mixed at the ratio of 1:9, 5:5, or 9:1. 150 µl of the mixed cells were inoculated onto the wells of the chitin sheet coated 96-well plate, and incubated at 25°C for more than 80 h. The growth profiles of the exploiter and the degrader were evaluated by measuring fluorescence intensity and $OD_{600}$ at 30-minute intervals with a plate reader (Tecan Spark).

For SYBR-gold staining, 100 ul samples were taken from cultures and allowed to incubate for 15 minutes in the dark, after which they were measured with a plate reader (Tecan Spark) using a SYBR-green filter set.

Genome databases and functional annotation

Genome bundles were downloaded from the NCBI Refseq FTP website on July 13, 2019 (*3*). Only the latest versions of complete genomes were downloaded, resulting in 13,737 genomes. We then removed redundant genomes from our dataset by clustering genomes that had >99.9% ANI over 90% of their genome (*4*), keeping the longest genome from each cluster, resulting in 8753 non-redundant genomes (*5*, *6*). The phylogenetic tree, as well as the classification of each genome, was obtained using GTDB-tk (*7*). For all downstream analyses, a concatenated file was created where only CDSs and their corresponding translated proteins were retained, with pseudogenes (as annotated by NCBI) discarded.

Chitinases (EC.3.2.1.14) were identified using a slightly modified dbCAN annotation workflow (*8*). The modified workflow consisted of identifying proteins that significantly aligned to the dbCAN HMMs (e-value < 10-10), and contained conserved k-mers, as identified by the dbCAN hotpep program (with hits-cutoff set to 1, and freq cutoff set to 0.2. These parameters were found to be the best performing on bacterial genomes (*8*)). Finally, hits were filtered to only contain GH18 and GH19, as these are the most prevalent and specific chitinase domains found in the CAZy database.

eggnog-mapper v2 was used to functionally annotate proteins, with parameters --go_evidence non-electronic --target_orthologs all --seed_ortholog_evalue 0.001 --seed_ortholog_score 60 (*9*). For KEGG annotation of co-evolving genes (Supplementary Figure S1), unknown function genes were genes that either had no KEGG orthology hit and no description or were not found in the eggnog output file (had no hits to seed orthologues). Mobile genes were genes that had one of the following terms in their description: phage, tail, capsid, transposon, transposase, conjugation, or insertion element. Other functions were extracted from the KEGG pathway annotations of genes. The final distribution of functions of co-evolving genes was calculated as follows: the probability of finding an annotation in a given gene was calculated, and then the sum of each annotation across all genes is the final number reported in table S1.

Co-evolution and genome proximity

All 49,638,395 genes from the 13,373 complete genomes were clustered into gene families based on homology of their protein products using mmseqs2, with command-line options --cov-mode 0 --min-seq-id 0.5 -c 0.8 -s 7 --max-seqs 300 --cluster-mode 1 (*10*). This means that in order to be clustered genes must be at least 50% identical over 80% of the length of the longer protein product. After clustering, which resulted in gene 3,237,392 families, we determined the copy number of each gene family in each genome. To determine gain / loss events, we used the Castor R package to reconstruct the ancestral copy-numbers of each gene family using a maximum-likelihood approach, given the extent genome's copy number, and the GTDB-tk generated species tree, by using the function asr_squared_change_parsimony (*11*). Changes were then calculated for each edge in the tree and every gene family as the difference between the ancestral copy number and the descending node's copy number, and rounding to the nearest integer.

Co-evolution was defined as similarities in gene gain/loss events (*12*, *13*). Similarities in evolutionary gain/loss events (after correcting for changes in total genome sizes, as done in (*12*)) were computed using the cosine similarity metric. As high similarity could be due to random chance if the observed number of changes is too small, we only considered correlations between chitinase and genes that had at least 3 changes on the phylogeny, and were found in at least 5 genomes (*13*). This reduced the number of genes analyzed to 270,461 (1,889,229 gene families were only found in a single genome). We utilized a randomization test to assess the significance of the observed similarities with chitinase. For each gene, we identified the subtree in which it exists, and verified that chitinase also has >2 changes in that subtree. We then generated 104 random vectors from the entries of the change matrix (edge x gene) in that specific subtree, by randomly permuting the entries of each edge across different genes, while only sampling from genes that had >2 changes in that subtree. This process was performed independently for each edge in the subtree. The p-value of the correlation was the fraction of random similarities that showed a higher similarity than the one observed for the chitinase-gene pair. In cases where the distribution of a gene was not monophyletic, the common ancestor of all genomes carrying the gene was used to define the subtree in which it exists.

To detect genes that co-localize frequently with chitinases, we first created a list of all protein sequences that are proximal to chitinases. In any genome that contained a chitinase, we extracted the genes that were ≤ 5 genes away from a chitinase. If a genome contained more than

1 chitinase, this procedure was performed for each chitinase in the genome. Each one of these proteins belonged to a gene-family, as described above. This allowed us to determine all gene-families that appear next to chitinases at least once. For each gene family, we calculated the probability, P, of being proximal to a chitinase as the number of genomes in which it is proximal to a chitinase, divided by the total number of genomes it is found in. Following (*14*), we set a relaxed threshold of $P \leq 0.1$ to consider a gene as co-localized with chitinases.

The conditional probability histograms shown in figure 1D were computed as follows: For each of the 3576 accessory genes, we found the most recent common ancestor (MRCA) of all genomes containing that gene. The MRCA was used to extract a subtree descending from that MRCA node, over which the analysis for each gene was carried out. The probability of losing chitinases while retaining the accessory gene was the ratio between 1) the sum of edges over which chitinase copy number went from some positive number to zero, while the copy number of the accessory gene remained positive (e.g, parent node had 1 chitinase and 1 copy of the coevolving gene, while child node had 0 chitinases and 1 copy of the coevolving gene), and 2) the sum of edges over which the copy number of the accessory gene remained positive. The second conditional histogram was computed similarly, with the categories switched.

To detect clusters of co-evolving genes, we used weighted minhash (*15*) to quickly find groups of similar vectors among all genes that co-evolve with chitinase, using a low threshold (0.3) for similarity. Due to the low threshold and probabilistic nature of the minhash clustering algorithm, we then broke up clusters identified using the weighted minhash, as these clusters may contain illegitimately clustered vectors. This was done as described above while skipping the significance testing step. This resulted in a network of similarities between genes, where nodes in the network are genes, and edges are similarities. To detect communities in the graph, we used the Louvain clustering method as implemented in the igraph R package.

Determination of $E_{chi}$ and isolate ecological role classification

An elastic-net linear model was trained on all non-redundant publicly available complete genomes, with chitin-associated gene content clusters as features, and actual chitinase counts as the response. The chitin-associated gene content of each genome was determined based on the protein clustering performed in the previous segment. Each genome was represented by a binary vector, where each entry represented a cluster of co-evolving genes. Entries were 1 if the genome contained more than 30% of genes belonging to that cluster, and 0 otherwise. These vectors, together with a vector of associated chitinase copy numbers in each genome were fed to an elastic-net regression, as implemented in the cv.glmnet function from the glmnet R package. The parameters used were nfold = 5, alpha = 0.2, nlambda = 1e3. The model was trained on each phylum separately (except for proteobacteria, which were split into alpha-, and gamma-proteobacteria), and on the entire combined dataset. The statistics of each model in terms of mean-square-error and cross-validated $R^2$ are found in supplementary table S4. The values of $E_{chi}$ were rounded to the nearest integer and genomes were classified according to the following scheme: genomes with $O_{chi} \geq 1$ and $E_{chi} \geq 1$ were classified as degraders. Genomes with $O_{chi} = 0$ & $E_{chi} \geq 1$ were classified as exploiters. Genomes with $O_{chi} = 0$ and $E_{chi} = 0$ were classified as scavengers. Additionally, we reasoned that organisms that have chitinases but are not retaining their chitinase-associated genes were not ecological primary degraders of chitin, and so genomes with $O_{chi} \geq 1$ & $E_{chi} = 0$ were classified as scavengers. Based on the relative rate of chitinase loss and chitinase-associated gene retention, compared to the opposite scenario, we postulate that such genomes evolved from chitinase containing ancestors, that lost their chitinases but retained many of their chitinase associated genes.

Genome-scale metabolic model construction was performed with CarveMe (*16*). Genes were predicted using prodigal (*17*), and functional annotation was performed in CarveMe with gap-filling on MBL media with pyruvate as the sole carbon source, as most isolates grew in that condition. Flux-balance analysis was performed using CobraPy with default parameters (*18*). Growth on GlcNAc was defined as a non-zero simulated biomass yield on MBL + 20 [mM] GlcNAc. Degraders were defined as genomes predicted to grow on GlcNAc and that possess a chitinase, exploiters were predicted to grow on GlcNAc but have no chitinases, and scavengers were genomes predicted to not grow on GlcNAc.

<u>Metagenomic 16S rRNA dynamics and early colonization rate calculation</u>

For Figure 3, the 16S rRNA sequence of each genome was determined by first amplifying it using the 27F (5-AGAGTTTGATCMTGGCTCAG-3) + 1492R (5-GGTTACCTTGTTACGACTT-3) universal bacterial primers(*19*, *20*), followed by sanger sequencing. Our strain collection was isolated from two experiments previously performed in the lab, which studied community succession on model particulate matter using natural sea-water communities (*19*, *20*). We mapped the 16S sequences of our isolates back to the metagenomic 16S trajectories (PRJNA319196, PRJNA478695) using blast, only keeping perfect matches (*21*). Each sequence was only mapped to the experiment from which it was isolated. The read counts were transformed into relative frequencies and only entries with a relative abundance > $10^{-3}$ were retained for downstream analysis. Each experiment was originally performed in triplicate, and we filtered the data for each isolate by only considering repeats that were coherent with each other, with a threshold of a Pearson correlation of 0.3 between repeats. The early colonization window was defined as the time window between 0 - 48 [h] for data derived from (*19*) or 0-60 [h] for data derived from (*20*), as the dynamics in this experiment were slower. The early colonization rate was defined as the average rate of increase in relative frequency between the lowest and highest points in the early colonization window. This rate was calculated for each repeat, and the average rate was returned for each isolate.

<u>Synthetic community experiment</u>

For the data presented in Figure 4, all isolates were pregrown in individual wells of a 96-well plate containing MB media for 48 hours. The individual pre-cultures were then combined in equal volumes, washed in MBL salts, and inoculated into deep-well 96-well plates containing MBL with 40 mg colloidal chitin as the sole carbon source. In each dilution cycle, communities were allowed to grow for 84 hours, and then diluted into fresh media with the appropriate dilution factor (1/10,1/100, or 1/1000). To determine community dynamics through time, DNA was extracted using the beckman-coulter DNAdvance kit, and sequenced using the EMP 16S amplicon protocol (using 515F (parada)-806R(apprill) primers) at the Environmental Sample Preparation and Sequencing Facility (ESPSF) which is located in Argonne National Labs.

<u>Consumer-resource model</u>

Co-cultures were modeled using a simple consumer resource model with the following equations:

$$\frac{dChitin}{dt} = -\alpha * Degrader * Chitin$$
$$\frac{dGlcNAc}{dt} = \alpha * Degrader * Chitin - GlcNAc * \left(\gamma_{deg} * Degrader + \gamma_{\exp} * Exploiter\right)$$
$$\frac{dDegrader}{dt} = Degrader * \left(\gamma_{deg} * GlcNAc - \lambda\right)$$
$$\frac{dExploiter}{dt} = Exploiter * \left(\gamma_{\exp} * GlcNAc - \lambda\right)$$

With α representing the hydrolysis rate, γ representing the uptake rate (assuming 100% efficiency of biomass conversion for simplicity) and λ representing a mortality rate. The model was solved numerically using the Julia package “DifferentialEquations” with default parameters. The model parameters were set to $\alpha = 2*10^{-4}$, $\gamma_{degrader} = 10^{-2}$, $\lambda = 10^{-2}$. Initial conditions were $Chitin_0 = 10^3$, $GlcNAc_0 = 0$, $degrader_0 = 10$, and $exploiter_0 = 10$. The model was simulated for 150 time integration steps. Different exploiter advantage ratios ( $\gamma_{exploiter} / \gamma_{degrader}$) were tested and are plotted in Supplementary Figure S12.

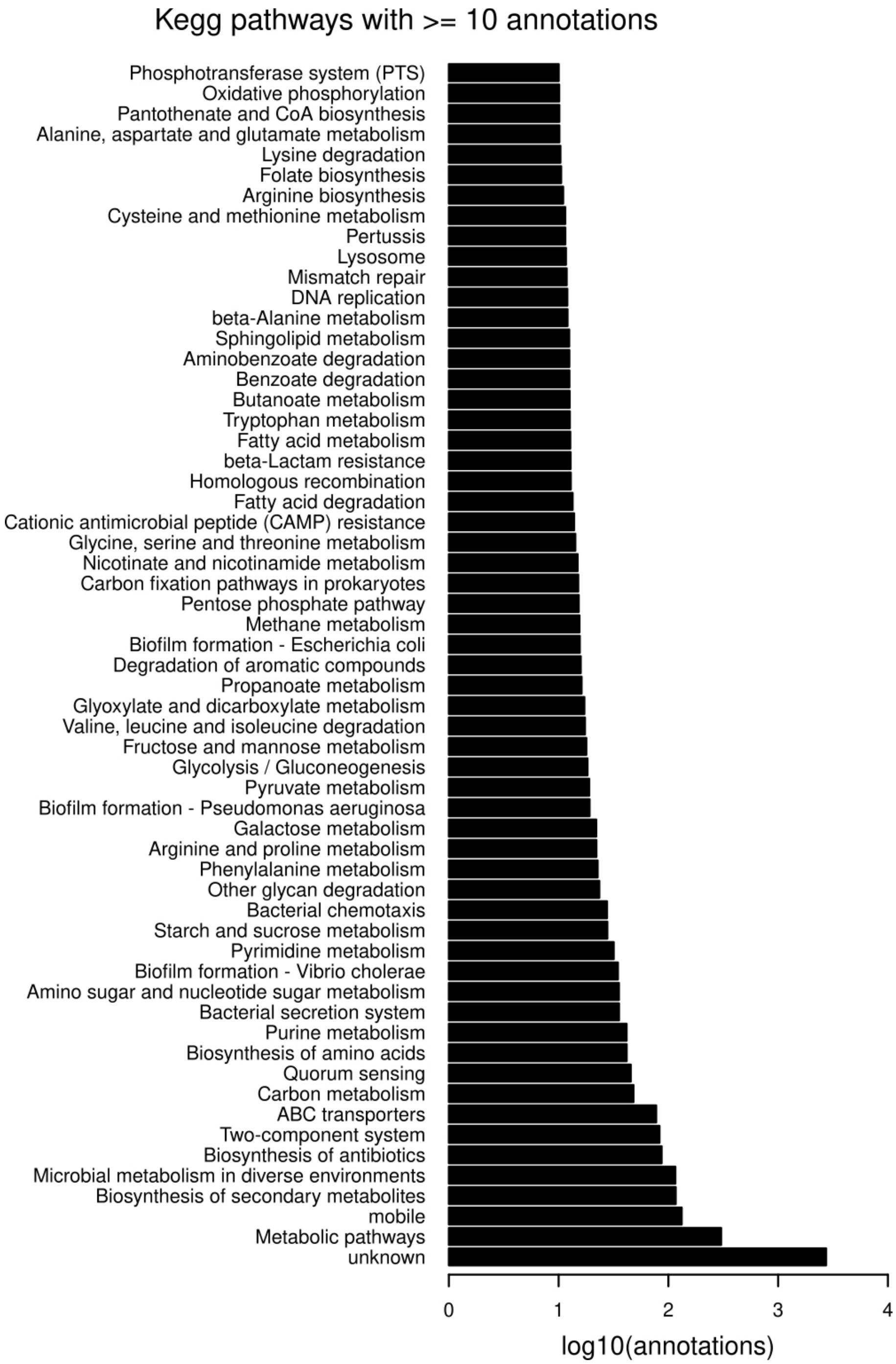


**Fig. S1. Distribution of KEGG annotations in chitinase linked genes.** Annotations were inferred using eggnog-mapper. Annotations are weighted in each gene, so that the total number of annotations in each gene is 1. The total number of annotations reported here is the sum of all weighted annotations across all genes. Mobile genes are genes that had one of the following terms in their description: phage, tail, capsid, transposon, transposase, conjugation, or insertion element. Unknown function genes either had no match to an eggnog-mapper entry, had no KEGG pathway annotation, or contained the term "unknown function" in the description field.

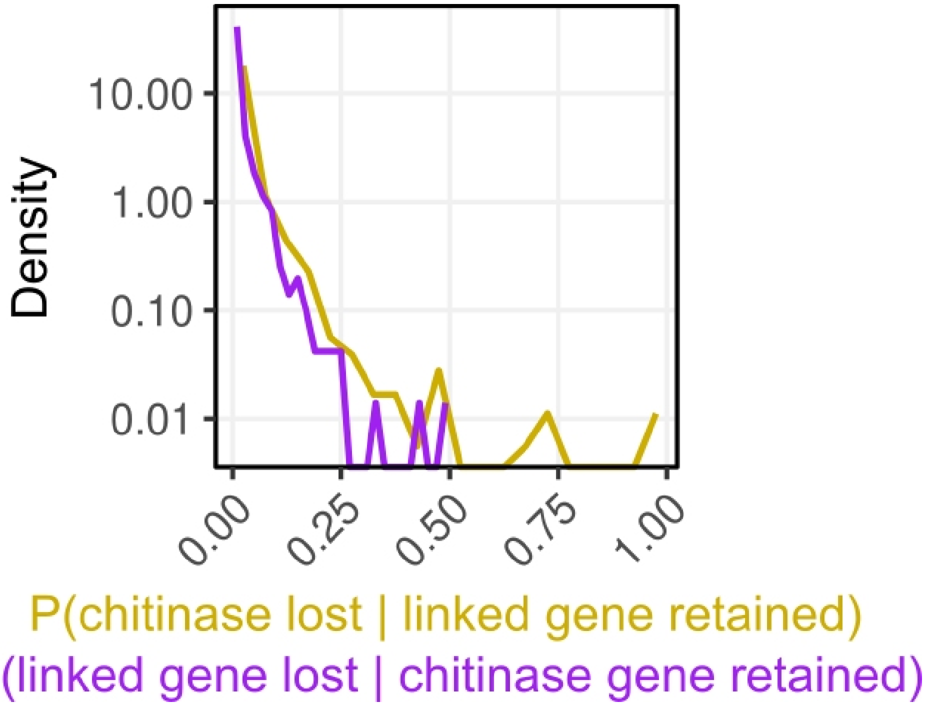


**Fig. S2. Density of conditional gene loss probabilities, given that the other gene category is retained.** The probability was calculated for each gene family across all edges of the relevant sub-tree over which the gene family exists. Unlike Fig. 1D, here the density is plotted with a logarithmic y axis and linear x axis, allowing the comparison of the densities at 0 conditional probability (0 x-axis values).

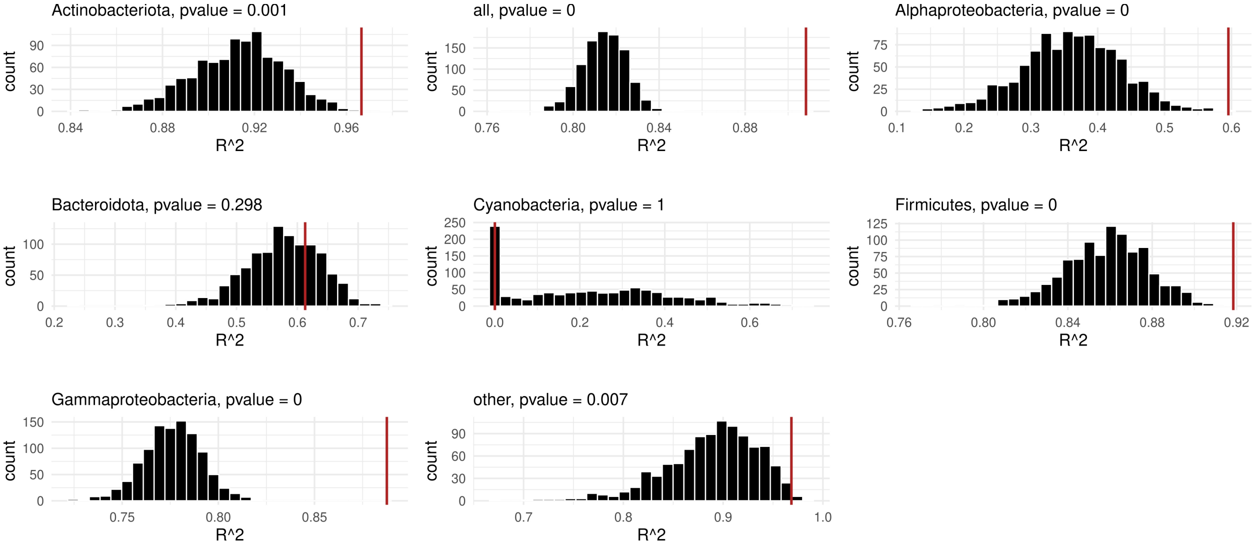


**Fig. S3. Chitinase linked genes are better than random sets of genes at predicting genomic chitinase copy number.** For each phylum/class the ability of our elastic-net linear regression to predict chitinase copy number, in terms of the $R^2$ of the regression, was compared to 1000 models trained with the same parameters on an equal number of randomly selected genes. The distribution of $R^2$ values of these random gene-based elastic-net linear models is plotted for each phylum/class, where the indicated p-value is the proportion of cases that a random gene elastic-net model had a higher $R^2$ than the chitinase linked genes-based elastic-net linear model. In cases where the p-values were insignificant, the successful random sets of genes either contained a large amount of co-evolving genes or contained many genes that were negatively linked to chitinases (data not shown). These negatively linked genes also contain valuable information about chitinase copy-number, but their functional significance in the context of chitin degradation is unclear.

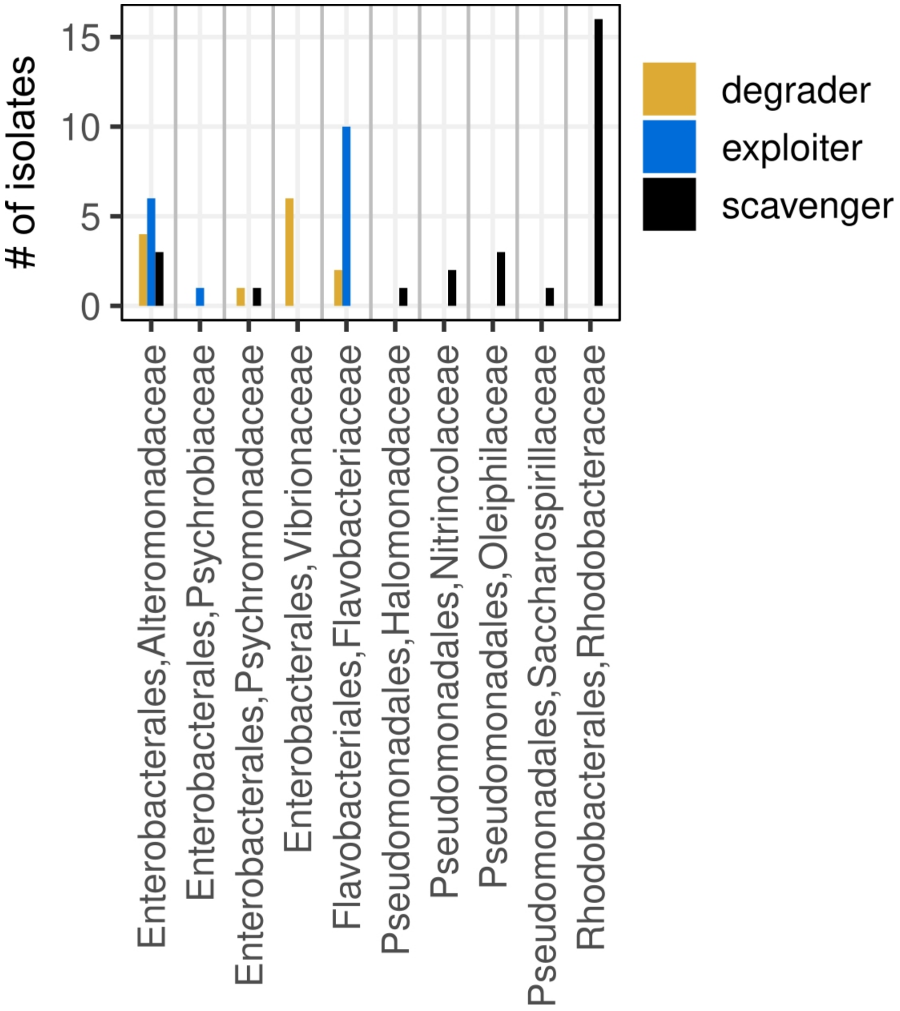


**Fig. S4. Predicted classes of 63 marine isolates (out of which 57 were phenotyped).** For each bacterial order in our collection (as determined by GTDB-tk), we predicted the expected number of chitinases ($E_{chi}$) based on the chitinase linked gene content of the isolate's genome, and compared it to the observed number of chitinases. All ecological roles were found in multiple orders and families, and for some orders contained multiple ecological roles, suggesting that ecological roles evolve faster than, and are unlinked from phylogeny at the order level.

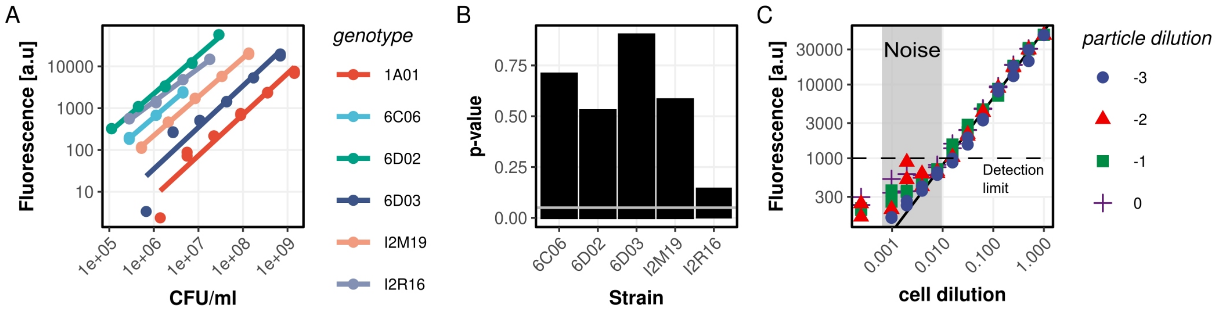


**Fig. S5. Accurate quantification of bacterial cell counts on chitin using SYBR-gold.** (A) fluorescence of individual strains as a function of cell density (CFU/ml). Cells were grown in marine broth 2216 to saturation and were then serially diluted and stained with SYBR-gold for 15 minutes, after which their fluorescence was determined. (B) P-values of the interaction term between strain identity and cell concentration for the regression lines of the data presented in panel A. No interaction term is significant (all p-values > 0.1), indicating that cell concentration influences fluorescence in a similar way across all tested strains. (C) Strain vibrio1A01 was stained with SYBR-gold in the presence of various concentrations of colloidal chitin. Particle dilutions are relative to the standard concentration used for assaying growth, which is 2 [g/l]. Cell concentrations are indicated as the dilution factor relative to the saturation cell density, which is ~$10^9$ [cells/ml]. The shaded region indicates the dilution factor where particle fluorescence overwhelmed cellular DNA fluorescence. The dashed line indicates our detection threshold, which was used in Figure 2A to define growth.

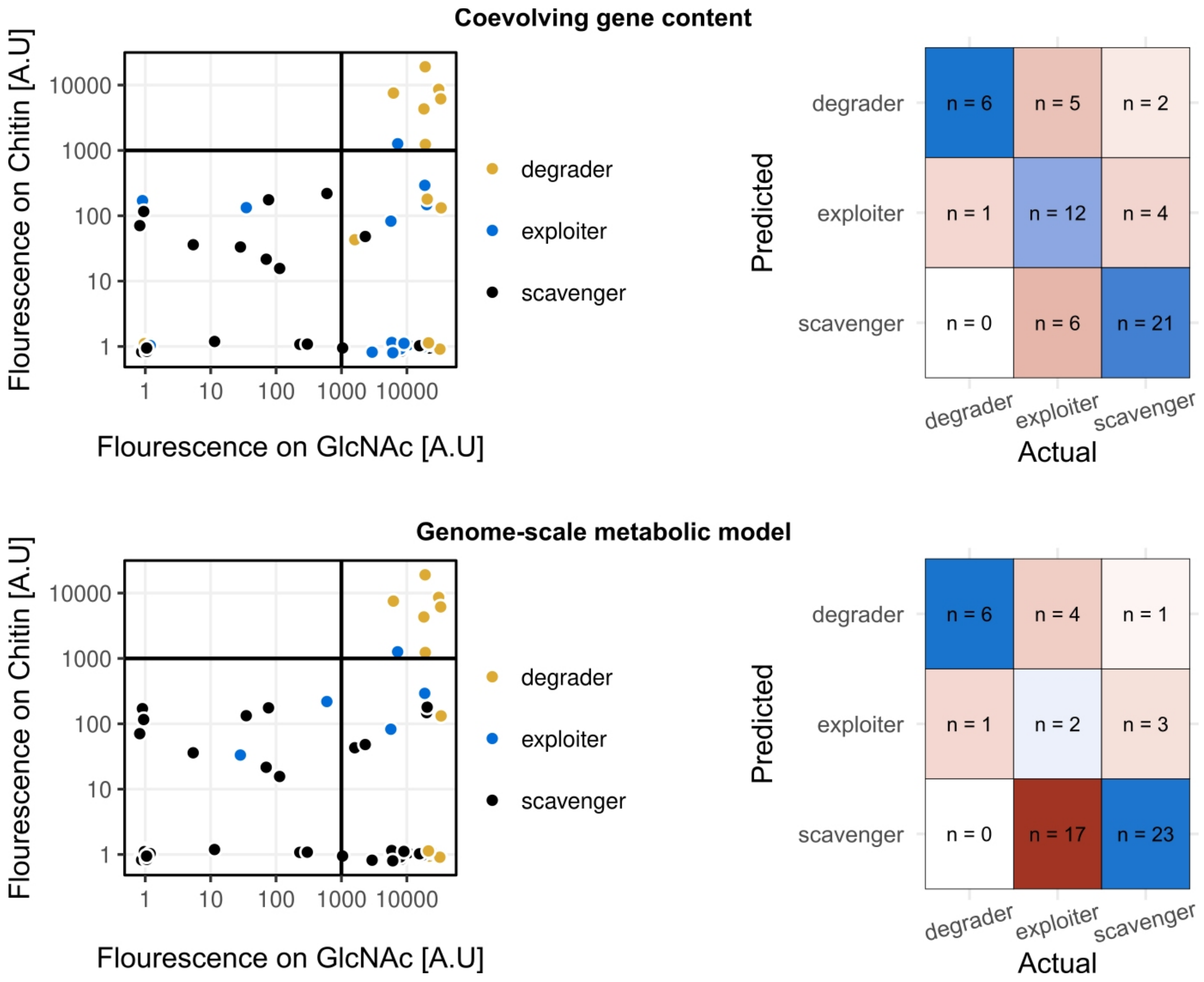


**Fig. S6. Confusion matrix for the two models.** For each model, the underlying data, which is the same data as presented in Fig. 2A, is presented on the left, where the points are colored according to the predictions of the indicated model. The confusion matrix that compares the predicted labels to observed labels is presented on the right. The major differences between the models are in their ability to identify exploiters.

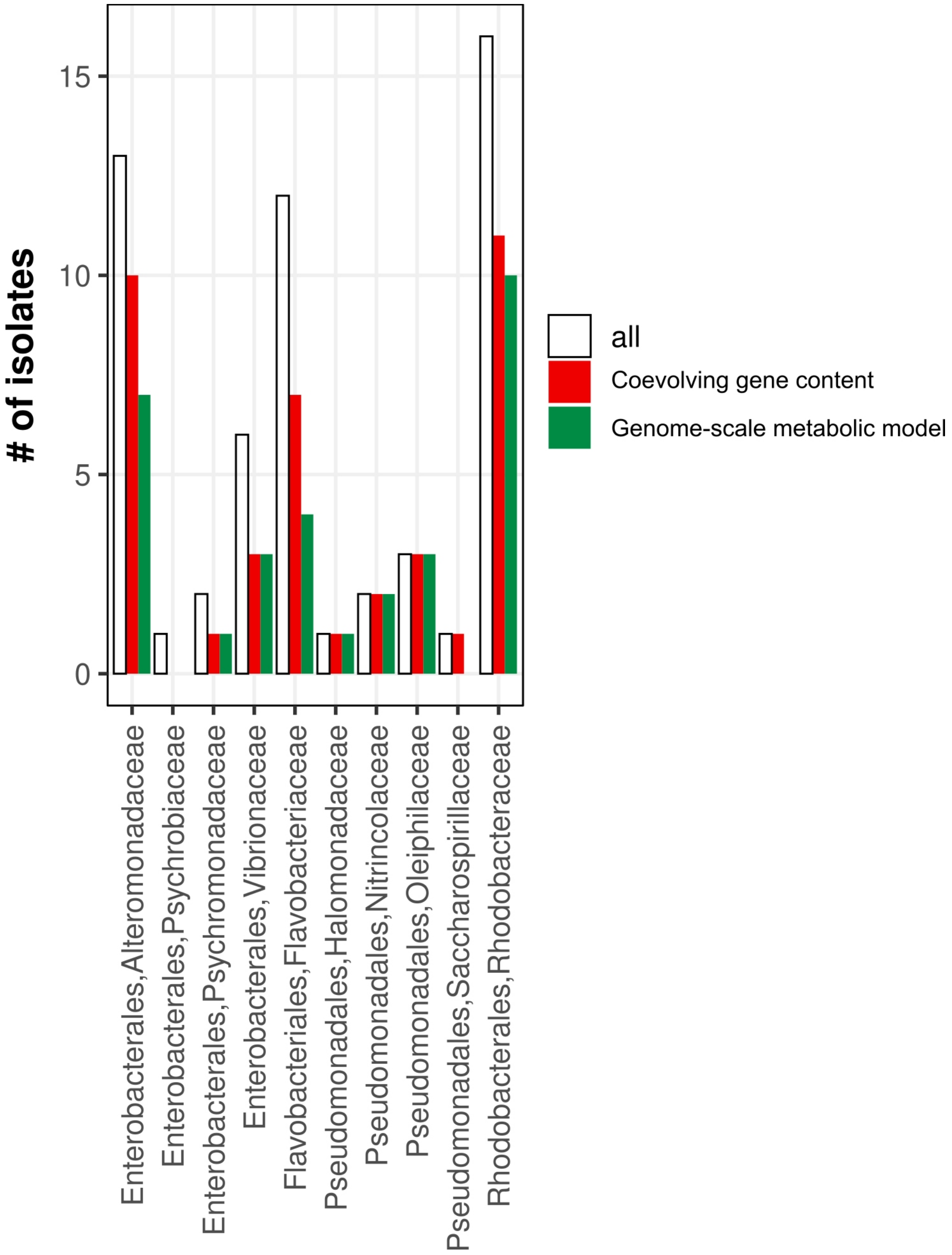


**Fig. S7. Genome-scale metabolic models perform worse in understudied taxa, compared to our (nearly) annotation free coevolving gene model.** The total number of isolates in each order are shown in white, and the number of isolates from that order correctly predicted by each type of model is indicated by the colored bars. A correct prediction is defined as in the main text, and the correct predictions of all ecological classes are amalgamated for the quantification in this figure.

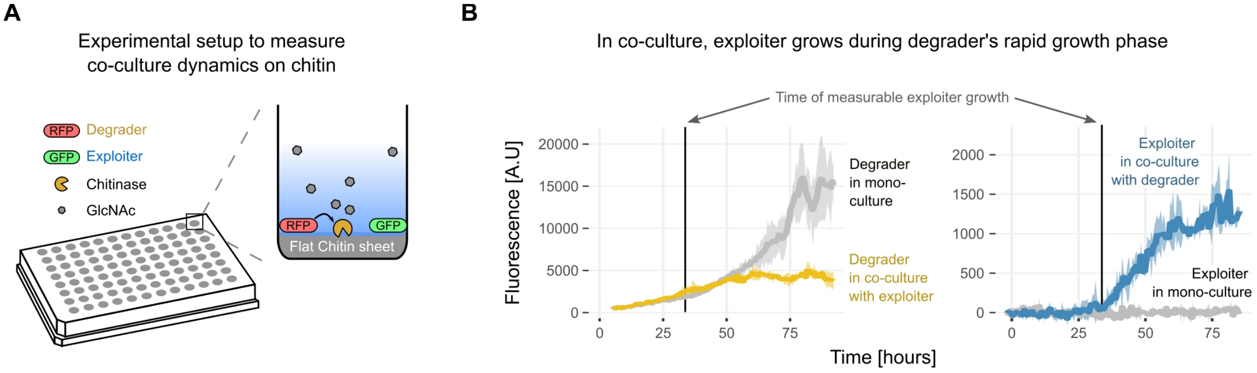


**Fig. S8. Dynamics of an exploiter–degrader co-culture. (A)** Illustration of the flat-chitin experimental setup used to measure co-culture dynamics. Strains are marked by fluorescent proteins expressed from a plasmid encoded constitutively expressed construct (methods) **(B)** Degrader-Exploiter co-culture reveals the consequence of exploitation among co-inhabiting environmental isolates. The Y-axis displays the fluorescence intensity, which is highly correlated by OD (Supplementary Figure S12). Grey lines represent mono-culture readings, while colored lines represent readings from co-cultures. Shaded regions represent standard deviations from the mean from three replicates**.** Vertical lines represent the earliest time where exploiter growth was observed when co-cultured with a degrader.

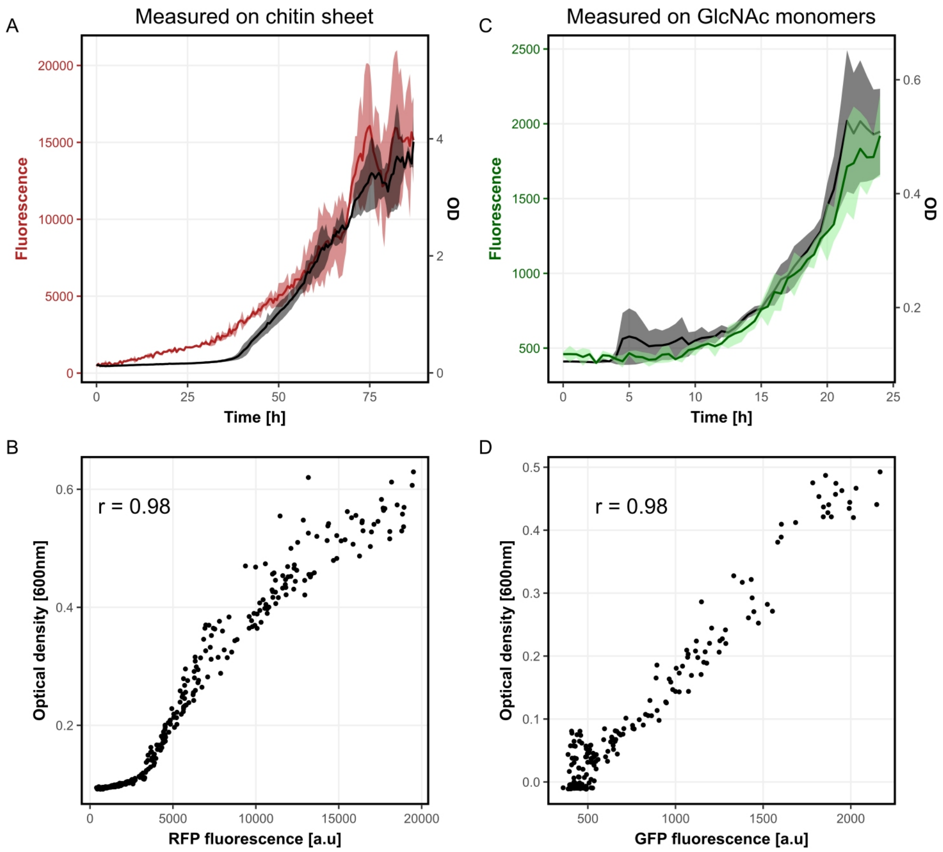


**Fig. S9. Plasmid encoded fluorescence correlates with OD.** (A,C) Fluorescence / Optical density (600nm, OD) as a function of time for strain vibrio1A01-DsRed (A) and strain alteroA3R04-eGFP (C). Red represents DsRed fluorescence, green represents eGFP fluorescence, and black represents OD. Experiments were conducted in triplicate, and the lines present the mean, with the shaded region representing the standard deviation. The DsRed reporter has a higher signal-to-noise compared with the eGFP reporter, resulting in higher background subtracted values, as well as the ability to detect an increase in fluorescence before an increase in OD. 1A01-DsRed measurements were performed with the chitin sheet as a sole carbon source, as in Fig. 2. alteroA3R04 is unable to grow in monoculture on the chitin sheet, so measurements were performed using the same protocol, but with 20 [mM] N-Acetylglucosamine added, resulting in faster dynamics. (B,D) correlation between fluorescence and optical density for vibrio1A01-DsRed (B) and alteroA3R04 (D). Values are taken from A and D respectively.

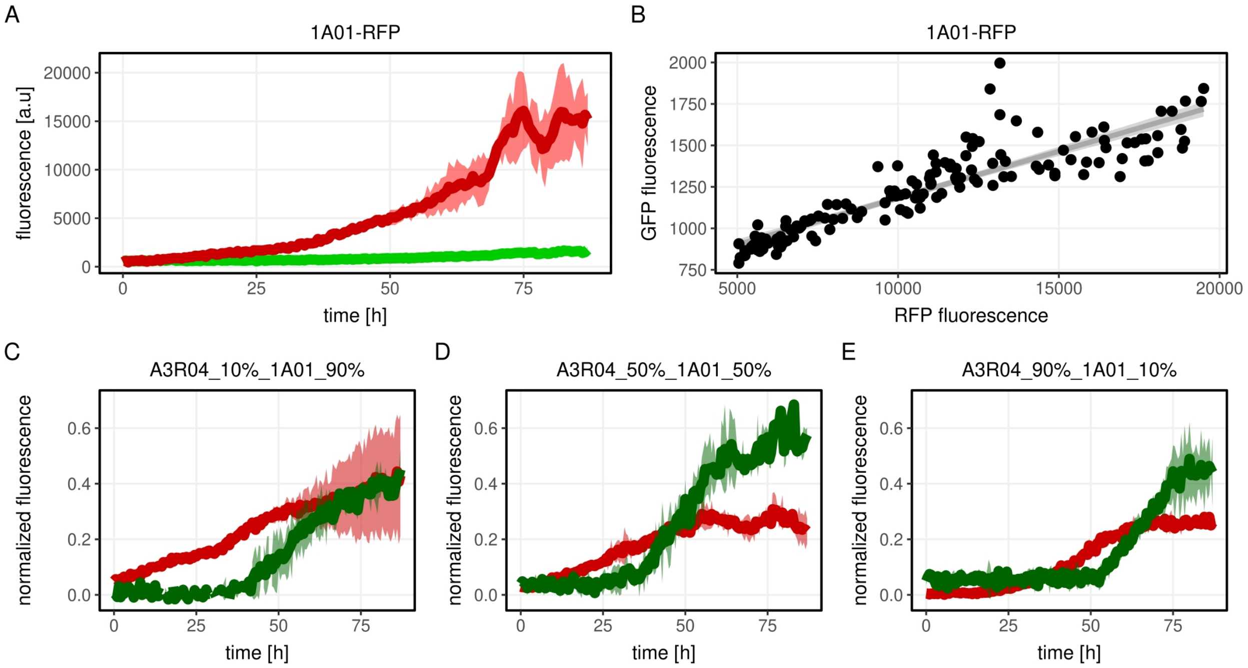


**Fig. S10. Using plasmid-based fluorescence to measure co-culture dynamics on chitin sheets.** (A,C) GFP and YFP fluorescence as a function of time for vibrio1A01-DsRed (A), and alteroA3R04-eGFO (C). Data is reproduced from experiments presented in Fig. S7, with the measurements of the additional fluorescent channel presented. Red lines represent DsRed fluorescence, Green lines represent eGFP fluorescence, and shaded regions represent standard deviations around the mean, taken from at least 2 indipendent replicates. (B,D) Correlation between DsRed and eGFP fluorescence for vibrio1A01-DsRed (B), and alteroA3R04-eGFP (D). The fit in B was used to correct eGFP values in co-cultures presented in panels E-G and in Fig. 2 in the main text. (E-G) co-culture dynamics of vibrio1A01-DsRed and alteroA3R04-eGFP when grown with a flat chitin sheet as the sole carbon source. To show the effect of the initial ratio of the two strains on the outcome of the co-culture, values were normalized such that the minimal DsRed/eGFP fluorescence value across all experiments was set to 0 and the maximal value across all experiments was set to 1. The data in F is used in Fig. 2 in the main text.

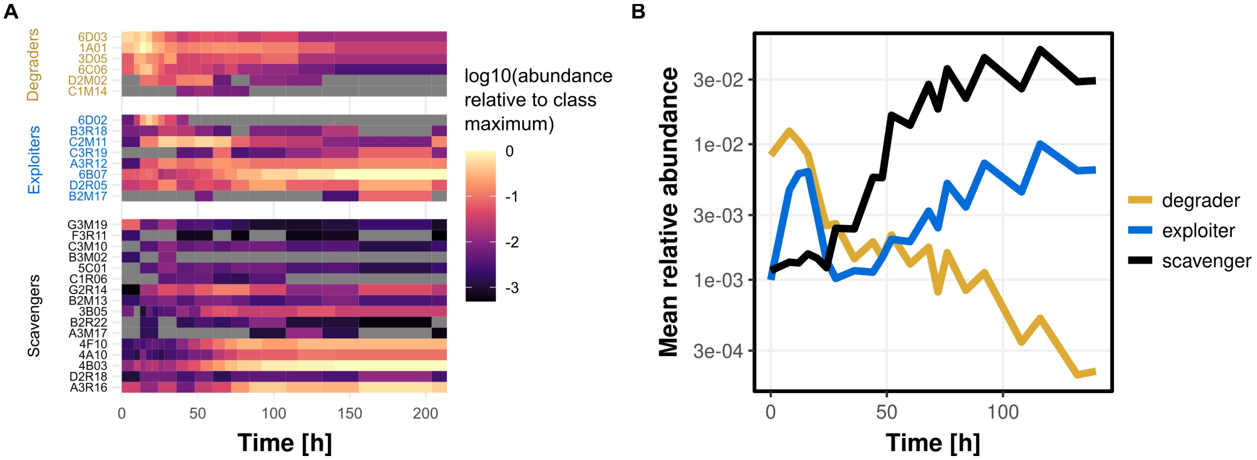


**Fig. S11. Dynamics of strains belonging to different ecological classes in a highly complex natural seawater community.** (A) dynamics of individual strains. Raw relative abundances were calculated as described in the methods section. Tiles are colored according to the relative abundance, divided by the maximal relative abundance of any isolate at any time point in the given ecological class. Grey tiles are below detection limit (relative frequency of $10^{-4}$). (B) Average relative abundance (not normalized as in A) of each ecological class, as a function of time.

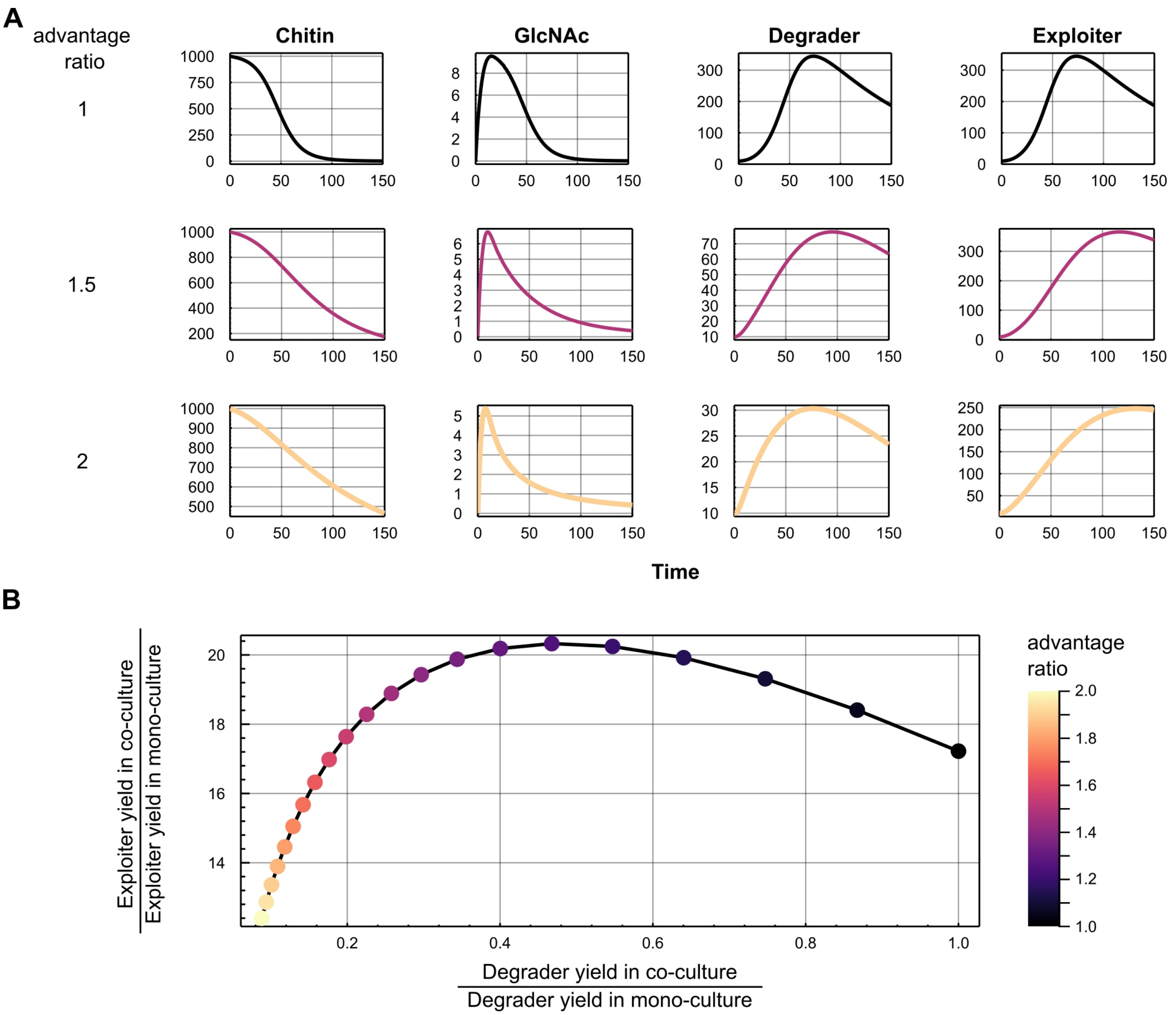


**Fig. S12. Simple consumer-resource model captures co-culture yield patterns.** (A) community and resource dynamics for 3 exploiter advantage ratios. The exploiter advantage ratio is defined as $\gamma_{exploiter} / \gamma_{degrader}$. The model parameters were set to $\alpha = 2*10^{-4}$, $\gamma_{degrader} = 10^{-2}$, $\lambda = 10^{-2}$. Initial conditions were $Chitin_0 = 10^3$, $GlcNAc_0 = 0$, $degrader_0 = 10$, and $exploiter_0 = 10$. The model was simulated for 150 time steps. (B) maximal yield of simulated co-cultures as a function of the exploiter advantage ratio. The axes show the ratio between the yield in co-culture and the yield in monoculture for the degrader (x-axis) and exploiter (y-axis). The exploiter monoculture was assumed to go through 1 doubling, but this choice only modulates the numbers on the y-axis, and does not change the qualitative outcome of the figure. The axes directly correspond to the axes in Figure 2C of the main text.

**Table S1.**

Number of genomes analyzed in each phylum or class, and the index by which the phylum is indicated in Fig. 1A.

| Phylum or class | # of genomes | number in figure 1A |
|---|---|---|
| Gammaproteobacteria | 3550 | 1 |
| Firmicutes | 2194 | 4 |
| Actinobacteriota | 928 | 6 |
| Alphaproteobacteria | 711 | 2 |
| other | 492 | none (grey edges) |
| Bacteroidota | 467 | 7 |
| Campylobacterota | 295 | 3 |
| Cyanobacteria | 115 | 5 |

**Table S2.**

Elastic-net regression summary statistics. For each phylum, the number of genomes analyzed, number of input features (clusters of chitinase linked genes that are found in at least one genome in that phylum or class), number of variables used by the elastic-net regression model, mean-square error of the model, and 5-fold cross-validated $R^2$. Values were obtained from the output of the cv.glmnet function in the glmnet R package.

| **Phylum or class** | **# of genomes** | **# of input features** | **# variables used** | **Mean square error** | **$R^2$** |
|---|---|---|---|---|---|
| all | 8752 | 1905 | 875 | 0.27 | 0.91 |
| Gammaproteobacteria | 3550 | 887 | 460 | 0.3 | 0.89 |
| Firmicutes | 2194 | 569 | 234 | 0.18 | 0.92 |
| Actinobacteriota | 928 | 518 | 206 | 0.28 | 0.97 |
| Alphaproteobacteria | 711 | 202 | 37 | 0.06 | 0.6 |
| other | 492 | 265 | 149 | 0.09 | 0.97 |
| Bacteroidota | 467 | 140 | 55 | 0.47 | 0.61 |
| Campylobacterota | 295 | 21 | 0 | 0.03 | 0 |
| Cyanobacteria | 115 | 39 | 0 | 0.26 | 0 |

**Table S3. Isolate phenotypes and model predictions.** The data underlying Fig 2A and Supplementary Figure S9 is given for each isolate in the columns labeled glcnac and chitin. The numbers represent the mean maximal fluorescence on each carbon source during 96 h of growth in monoculture. The phenotype is classification is derived from these measurements, where growth is considered as showing a fluorescence value > 1000. Degraders grow on both carbon sources, exploiters only on GlcNAc, and scavengers on neither. The predictions of both our co-evolution based model, and a CarveMe derived genome-scale metabolic model are indicated for each isolate. GTDB-tk derived phylogeny down to the genus level is also provided for each isolate

| strain | Phylum | Class | Order | Family | Genus | glcnac | chitin | phenotype | Co-evolving gene content | Genome-scale metabolic model |
|---|---|---|---|---|---|---|---|---|---|---|
| 1A01 | Proteobacteria | Gammaproteobacteria | Enterobacterales | Vibrionaceae | Vibrio | 30680.65 | 8595.6 | degrader | degrader | degrader |
| 3B05 | Proteobacteria | Gammaproteobacteria | Pseudomonadales | Nitrincolaceae | Neptunomonas | 112.48 | 14.49 | scavenger | scavenger | scavenger |
| 3D05 | Proteobacteria | Gammaproteobacteria | Enterobacterales | Alteromonadaceae | Pseudoalteromonas | 19161.67 | 1233 | degrader | degrader | degrader |
| 4A10 | Proteobacteria | Alphaproteobacteria | Rhodobacterales | Rhodobacteraceae | Epibacterium | 0 | 0 | scavenger | scavenger | scavenger |
| 4B03 | Proteobacteria | Gammaproteobacteria | Enterobacterales | Alteromonadaceae | Alteromonas | 0 | 69.54 | scavenger | scavenger | scavenger |
| 4F10 | Proteobacteria | Alphaproteobacteria | Rhodobacterales | Rhodobacteraceae | Pseudopelagicola | 0 | 0 | scavenger | scavenger | scavenger |
| 5C01 | Proteobacteria | Alphaproteobacteria | Rhodobacterales | Rhodobacteraceae | OMPR01 | 230 | 0 | scavenger | scavenger | scavenger |
| 6B07 | Bacteroidota | Bacteroidia | Flavobacteriales | Flavobacteriaceae | Maribacter | 10437.74 | 0 | exploiter | exploiter | scavenger |
| 6C06 | Proteobacteria | Gammaproteobacteria | Enterobacterales | Psychromonadaceae | Psychromonas | 6232.23 | 7542.26 | degrader | degrader | degrader |
| 6D02 | Proteobacteria | Gammaproteobacteria | Enterobacterales | Psychrobiaceae | Psychrobium | 7233.89 | 1263.15 | degrader | exploiter | exploiter |
| 6D03 | Proteobacteria | Gammaproteobacteria | Enterobacterales | Vibrionaceae | Vibrio | 19031.74 | 19044.56 | degrader | degrader | degrader |
| A2M03 | Bacteroidota | Bacteroidia | Flavobacteriales | Flavobacteriaceae | Zobellia | 1597.28 | 41.89 | exploiter | degrader | scavenger |
| A3M17 | Proteobacteria | Alphaproteobacteria | Rhodobacterales | Rhodobacteraceae | Ruegeria | 2312.22 | 47.17 | exploiter | scavenger | scavenger |
| A3R04 | Proteobacteria | Gammaproteobacteria | Enterobacterales | Alteromonadaceae | Alteromonas_B | 18035.39 | 0 | exploiter | exploiter | scavenger |
| A3R06 | Proteobacteria | Alphaproteobacteria | Rhodobacterales | Rhodobacteraceae | Tritonibacter | 8193.27 | 0 | exploiter | scavenger | scavenger |
| A3R12 | Proteobacteria | Gammaproteobacteria | Enterobacterales | Alteromonadaceae | Paraglaciecola | 0 | 0 | scavenger | exploiter | scavenger |
| A3R16 | Proteobacteria | Gammaproteobacteria | Enterobacterales | Alteromonadaceae | Paraglaciecola | 0 | 0 | scavenger | scavenger | exploiter |
| B2M06 | Bacteroidota | Bacteroidia | Flavobacteriales | Flavobacteriaceae | Cellulophaga | 7946.93 | 0 | exploiter | exploiter | scavenger |
| B2M13 | Proteobacteria | Gammaproteobacteria | Pseudomonadales | Halomonadaceae | Cobetia | 0 | 0 | scavenger | scavenger | scavenger |
| B2M17 | Bacteroidota | Bacteroidia | Flavobacteriales | Flavobacteriaceae | Polaribacter | 0 | 169.33 | scavenger | exploiter | scavenger |
| B2R09 | Proteobacteria | Alphaproteobacteria | Rhodobacterales | Rhodobacteraceae | Roseovarius | 0 | 0 | scavenger | scavenger | scavenger |
| B2R22 | Proteobacteria | Alphaproteobacteria | Rhodobacterales | Rhodobacteraceae | Octadecabacter | 0 | 0 | scavenger | scavenger | scavenger |
| B3M02 | Proteobacteria | Gammaproteobacteria | Enterobacterales | Psychromonadaceae | Psychromonas | 21837.89 | 0 | exploiter | scavenger | degrader |
| B3M08 | Proteobacteria | Alphaproteobacteria | Rhodobacterales | Rhodobacteraceae | Lentibacter | 5860 | 0 | exploiter | scavenger | scavenger |
| B3R10 | Proteobacteria | Gammaproteobacteria | Enterobacterales | Alteromonadaceae | Psychrosphaera | 20081.09 | 146.99 | exploiter | exploiter | scavenger |
| B3R18 | Bacteroidota | Bacteroidia | Flavobacteriales | Flavobacteriaceae | Zobellia | 23957.62 | 0 | exploiter | exploiter | scavenger |
| C1M14 | Proteobacteria | Gammaproteobacteria | Enterobacterales | Alteromonadaceae | Alteromonas_B | 20446.28 | 179.05 | exploiter | degrader | scavenger |
| C1R06 | Proteobacteria | Gammaproteobacteria | Pseudomonadales | Nitrincolaceae | Amphritea | 0 | 0 | scavenger | scavenger | scavenger |
| C2M11 | Proteobacteria | Gammaproteobacteria | Enterobacterales | Alteromonadaceae | Colwellia | 18780 | 291.28 | exploiter | exploiter | exploiter |
| C2R02 | Proteobacteria | Gammaproteobacteria | Enterobacterales | Alteromonadaceae | Pseudoalteromonas | 0 | 0 | scavenger | degrader | degrader |
| C2R09 | Proteobacteria | Alphaproteobacteria | Rhodobacterales | Rhodobacteraceae | Paracoccus | 27.33 | 32.11 | scavenger | scavenger | exploiter |
| C3M06 | Proteobacteria | Alphaproteobacteria | Rhodobacterales | Rhodobacteraceae | Salipiger | 69.63 | 20.8 | scavenger | scavenger | scavenger |
| C3M10 | Proteobacteria | Alphaproteobacteria | Rhodobacterales | Rhodobacteraceae | OMPR01 | 10.44 | 0 | scavenger | scavenger | scavenger |
| C3R12 | Proteobacteria | Gammaproteobacteria | Enterobacterales | Vibrionaceae | Vibrio | 31988.59 | 0 | exploiter | degrader | degrader |
| C3R17 | Bacteroidota | Bacteroidia | Flavobacteriales | Flavobacteriaceae | Zobellia | 2944.06 | 0 | exploiter | exploiter | scavenger |
| C3R19 | Bacteroidota | Bacteroidia | Flavobacteriales | Flavobacteriaceae | Maribacter | 17172.04 | 0 | exploiter | exploiter | scavenger |
| D2M02 | Proteobacteria | Gammaproteobacteria | Enterobacterales | Alteromonadaceae | Colwellia | 18222.93 | 4308.47 | degrader | degrader | degrader |

| | | | | | | | | | | |
|---|---|---|---|---|---|---|---|---|---|---|
| D2M19 | Proteobacteria | Gammaproteobacteria | Pseudomonadales | Oleiphilaceae | Marinobacter | 4.26 | 34.83 | scavenger | scavenger | scavenger |
| D2R04 | Proteobacteria | Alphaproteobacteria | Rhodobacterales | Rhodobacteraceae | Pacificibacter | 0 | 0 | scavenger | scavenger | scavenger |
| D2R05 | Proteobacteria | Gammaproteobacteria | Enterobacterales | Alteromonadaceae | Aliiglaciecola | 5693 | 81.48 | exploiter | exploiter | exploiter |
| D2R18 | Proteobacteria | Alphaproteobacteria | Rhodobacterales | Rhodobacteraceae | Yoonia | 15614.33 | 0 | exploiter | scavenger | scavenger |
| D3R19 | Bacteroidota | Bacteroidia | Flavobacteriales | Flavobacteriaceae | Winogradskyella | 34 | 132.15 | scavenger | exploiter | scavenger |
| E3M18 | Bacteroidota | Bacteroidia | Flavobacteriales | Flavobacteriaceae | Arenibacter | 5879.01 | 0 | exploiter | exploiter | scavenger |
| E3R01 | Bacteroidota | Bacteroidia | Flavobacteriales | Flavobacteriaceae | Tenacibaculum | 0 | 0 | scavenger | degrader | scavenger |
| E3R09 | Bacteroidota | Bacteroidia | Flavobacteriales | Flavobacteriaceae | Winogradskyella | 0 | 0 | scavenger | exploiter | scavenger |
| E3R11 | Proteobacteria | Gammaproteobacteria | Enterobacterales | Vibrionaceae | Vibrio | 21428 | 0 | exploiter | degrader | degrader |
| F2R14 | Proteobacteria | Alphaproteobacteria | Rhodobacterales | Rhodobacteraceae | Pacificibacter | 0 | 0 | scavenger | scavenger | scavenger |
| F3M07 | Proteobacteria | Gammaproteobacteria | Enterobacterales | Alteromonadaceae | Psychrosphaera | 0 | 115 | scavenger | scavenger | scavenger |
| F3R08 | Proteobacteria | Gammaproteobacteria | Pseudomonadales | Oleiphilaceae | Marinobacter | 299.88 | 0 | scavenger | scavenger | scavenger |
| F3R11 | Proteobacteria | Gammaproteobacteria | Pseudomonadales | Oleiphilaceae | Marinobacter | 75.73 | 174.48 | scavenger | scavenger | scavenger |
| G2M07 | Proteobacteria | Gammaproteobacteria | Pseudomonadales | Saccharospirillaceae | Reinekea | 597.24 | 218.18 | scavenger | scavenger | exploiter |
| G2R10 | Proteobacteria | Gammaproteobacteria | Enterobacterales | Vibrionaceae | Vibrio | 32911.67 | 6150 | degrader | degrader | degrader |
| G2R14 | Proteobacteria | Alphaproteobacteria | Rhodobacterales | Rhodobacteraceae | Pseudopelagicola | 0 | 0 | scavenger | scavenger | scavenger |
| G3M19 | Proteobacteria | Alphaproteobacteria | Rhodobacterales | Rhodobacteraceae | Celeribacter | 1039.31 | 0 | exploiter | scavenger | scavenger |
| I2M19 | Bacteroidota | Bacteroidia | Flavobacteriales | Flavobacteriaceae | Tamlana_A | 6087.67 | 0 | exploiter | exploiter | scavenger |
| I2R16 | Proteobacteria | Gammaproteobacteria | Enterobacterales | Alteromonadaceae | Psychrosphaera | 9020.68 | 0 | exploiter | exploiter | scavenger |
| I3M07 | Proteobacteria | Gammaproteobacteria | Enterobacterales | Vibrionaceae | Vibrio | 33400.33 | 131 | exploiter | degrader | degrader |

**Table S4. C.F.U counts of degrader/exploiter co-cultures and mono-cultures**. Isolates were grown in MBL minimal media with 2 g/l colloidal chitin as the sole carbon source, and their CFU counts were monitored before and after growth by plating. Co-cultured strains are differentiable by colony morphology.

| strain1 | strain2 | focal | type | time | replicate | cfu |
|---|---|---|---|---|---|---|
| 1A01 | 1A01 | 1A01 | mono | 0 | 1 | 1600000 |
| 6D03 | 6D03 | 6D03 | mono | 0 | 1 | 2200000 |
| C3R19 | C3R19 | C3R19 | mono | 0 | 1 | 1400000 |
| 6B07 | 6B07 | 6B07 | mono | 0 | 1 | 1800000 |
| I2M19 | I2M19 | I2M19 | mono | 0 | 1 | 400000 |
| 1A01 | 1A01 | 1A01 | mono | 72 | 1 | 1600000000 |
| 6D03 | 6D03 | 6D03 | mono | 72 | 1 | 400000000 |
| C3R19 | C3R19 | C3R19 | mono | 72 | 1 | 5000000 |
| 6B07 | 6B07 | 6B07 | mono | 72 | 1 | 3000000 |
| I2M19 | I2M19 | I2M19 | mono | 72 | 1 | 1000000 |
| 1A01 | 1A01 | 1A01 | mono | 72 | 2 | 1700000000 |
| 6D03 | 6D03 | 6D03 | mono | 72 | 2 | 400000000 |
| C3R19 | C3R19 | C3R19 | mono | 72 | 2 | 1000000 |
| 6B07 | 6B07 | 6B07 | mono | 72 | 2 | 10000000 |
| I2M19 | I2M19 | I2M19 | mono | 72 | 2 | 1200000 |
| 1A01 | C3R19 | 1A01 | co | 72 | 1 | 490000000 |
| 1A01 | C3R19 | 1A01 | co | 72 | 2 | 35000000 |
| 1A01 | C3R19 | C3R19 | co | 72 | 1 | 70000000 |
| 1A01 | C3R19 | C3R19 | co | 72 | 2 | 7000000 |
| 1A01 | 6B07 | 1A01 | co | 72 | 1 | 35000000 |
| 1A01 | 6B07 | 1A01 | co | 72 | 2 | 43000000 |
| 1A01 | 6B07 | 6B07 | co | 72 | 1 | 9000000 |
| 1A01 | 6B07 | 6B07 | co | 72 | 2 | 20000000 |
| 1A01 | I2M19 | 1A01 | co | 72 | 1 | 1020000000 |
| 1A01 | I2M19 | 1A01 | co | 72 | 2 | 410000000 |
| 1A01 | I2M19 | I2M19 | co | 72 | 1 | 30000000 |
| 1A01 | I2M19 | I2M19 | co | 72 | 2 | 20000000 |
| 6D03 | C3R19 | 6D03 | co | 72 | 1 | 55000000 |
| 6D03 | C3R19 | 6D03 | co | 72 | 2 | 72000000 |
| 6D03 | C3R19 | C3R19 | co | 72 | 1 | 40000000 |
| 6D03 | C3R19 | C3R19 | co | 72 | 2 | 23000000 |
| 6D03 | 6B07 | 6D03 | co | 72 | 1 | 14000000 |
| 6D03 | 6B07 | 6D03 | co | 72 | 2 | 36000000 |
| 6D03 | 6B07 | 6B07 | co | 72 | 1 | 15000000 |
| 6D03 | 6B07 | 6B07 | co | 72 | 2 | 40000000 |
| 6D03 | I2M19 | 6D03 | co | 72 | 1 | 24000000 |
| 6D03 | I2M19 | 6D03 | co | 72 | 2 | 83000000 |
| 6D03 | I2M19 | I2M19 | co | 72 | 1 | 14000000 |
| 6D03 | I2M19 | I2M19 | co | 72 | 2 | 8000000 |

**Data S1. (separate file)**

Description of the 8752 genomes used in the analysis, including: accession #, bioproject, biosample, wgs_master, refseq_category, taxid, species_taxid, organism_name, infraspecific_name, isolate, version_status, assembly_level, release_type, genome_rep, seq_rel_date, asm_name, submitter, gbrs_paired_asm, paired_asm_comp, ftp_path, excluded_from_refseq, relation_to_type_material.

**Data S2. (separate file)**
GTDB-tk summary table and accompanying phylogenetic tree.

**Data S3. (separate file)**

KEGG pathway annotation of chitinase linked gene families. For each gene family, the different eggnog nogs that members of this protein family map to are reported, with the percent of members that map to that nog given in parenthesis. The different KEGG pathways that members of the given protein family map to is given in the same format.

**Data S4. (separate file)**

Weighted KEGG pathway annotations of chitinase associated genes. Annotations are summed across all chitinase linked gene families, weighted by their relative abundance in the given gene family.

**Data S5. (separate file)**

Taxonomy, and phenotypic classification of 44 isolates used in the synthetic community experiment presented in Fig. 4.

1. A. K. Dunn, D. S. Millikan, D. M. Adin, J. L. Bose, E. V. Stabb, New rfp- and pES213-Derived Tools for Analyzing Symbiotic Vibrio fischeri Reveal Patterns of Infection and lux Expression In Situ. *Appl. Environ. Microbiol.* **72**, 802–810 (2006).
2. E. V. Stabb, E. G. Ruby, in *Methods in Enzymology* (Academic Press, 2002; http://www.sciencedirect.com/science/article/pii/S0076687902581064), vol. 358 of *Bacterial Pathogenesis Part C: Identification, Regulation, and Function of Virulence Factors*, pp. 413–426.
3. N. A. O'Leary, M. W. Wright, J. R. Brister, S. Ciufo, D. Haddad, R. McVeigh, B. Rajput, B. Robbertse, B. Smith-White, D. Ako-Adjei, A. Astashyn, A. Badretdin, Y. Bao, O. Blinkova, V. Brover, V. Chetvernin, J. Choi, E. Cox, O. Ermolaeva, C. M. Farrell, T. Goldfarb, T. Gupta, D. Haft, E. Hatcher, W. Hlavina, V. S. Joardar, V. K. Kodali, W. Li, D. Maglott, P. Masterson, K. M. McGarvey, M. R. Murphy, K. O'Neill, S. Pujar, S. H. Rangwala, D. Rausch, L. D. Riddick, C. Schoch, A. Shkeda, S. S. Storz, H. Sun, F. Thibaud-Nissen, I. Tolstoy, R. E. Tully, A. R. Vatsan, C. Wallin, D. Webb, W. Wu, M. J. Landrum, A. Kimchi, T. Tatusova, M. DiCuccio, P. Kitts, T. D. Murphy, K. D. Pruitt, Reference sequence (RefSeq) database at NCBI: current status, taxonomic expansion, and functional annotation. *Nucleic Acids Res.* **44**, D733–D745 (2016).
4. C. Jain, L. M. Rodriguez-R, A. M. Phillippy, K. T. Konstantinidis, S. Aluru, High throughput ANI analysis of 90K prokaryotic genomes reveals clear species boundaries. *Nat. Commun.* **9**, 1–8 (2018).
5. V. D. Blondel, J.-L. Guillaume, R. Lambiotte, E. Lefebvre, Fast unfolding of communities in large networks. *J. Stat. Mech. Theory Exp.* **2008**, P10008 (2008).
6. G. Csardi, T. Nepusz, The igraph software package for complex network research, 9.
7. P.-A. Chaumeil, A. J. Mussig, P. Hugenholtz, D. H. Parks, GTDB-Tk: a toolkit to classify genomes with the Genome Taxonomy Database. *Bioinformatics*. **36**, 1925–1927 (2020).
8. H. Zhang, T. Yohe, L. Huang, S. Entwistle, P. Wu, Z. Yang, P. K. Busk, Y. Xu, Y. Yin, dbCAN2: a meta server for automated carbohydrate-active enzyme annotation. *Nucleic Acids Res.* **46**, W95–W101 (2018).
9. J. Huerta-Cepas, K. Forslund, L. P. Coelho, D. Szklarczyk, L. J. Jensen, C. von Mering, P. Bork, Fast Genome-Wide Functional Annotation through Orthology Assignment by eggNOG-Mapper. *Mol. Biol. Evol.* **34**, 2115–2122 (2017).
10. M. Steinegger, J. Söding, MMseqs2 enables sensitive protein sequence searching for the analysis of massive data sets. *Nat. Biotechnol.* **35**, 1026–1028 (2017).
11. S. Louca, M. Doebeli, Efficient comparative phylogenetics on large trees. *Bioinformatics*. **34**, 1053–1055 (2018).
12. O. X. Cordero, B. Snel, P. Hogeweg, Coevolution of gene families in prokaryotes. *Genome Res.* **18**, 462–468 (2008).
13. O. Cohen, H. Ashkenazy, D. Burstein, T. Pupko, Uncovering the co-evolutionary network among prokaryotic genes. *Bioinformatics*. **28**, i389–i394 (2012).
14. L. Gao, H. Altae-Tran, F. Böhning, K. S. Makarova, M. Segel, J. L. Schmid-Burgk, J. Koob, Y. I. Wolf, E. V. Koonin, F. Zhang, Diverse enzymatic activities mediate antiviral immunity in prokaryotes. *Science*. **369**, 1077–1084 (2020).
